# Label-Free All-Electrical Tracking of Individual and Collective Cell Migration on a Megapixel CMOS Capacitance Sensor

**DOI:** 10.64898/2026.06.16.731623

**Authors:** Hyuntae Jeong, Pushkaraj Joshi, Yinshi Hu, Jiwon Kim, Anh Hoang Vu, Jacob K. Rosenstein, Ian Y. Wong

**Affiliations:** School of Engineering, Brown University. 184 Hope St Box D. Providence, Rhode Island, USA

## Abstract

Label-free tracking of adherent cell migration could enable important insights into biological processes such as tissue repair, inflammatory response, or cancer progression. Nevertheless, visualizing unlabeled animal cells using optical microscopy remains challenging due to low contrast as well as frequent changes in cell shape and number. A promising alternative uses electrical capacitance measurements, which are sensitive to cell adhesion to electrode surfaces. However, prior examples often utilized electrodes with areas larger than single cells, resulting in averaged readouts over multiple cells. Here, we demonstrate label-free, live-cell tracking using a capacitance sensor array with more than 1 million pixels on a 10 micron pitch across an area larger than 1 square centimeter. We show that single cell morphology can be clearly segmented, and then used to reconstruct migration and proliferation dynamics using optical flow. We further track the spreading of multicellular spheroids, revealing fast-moving peripheral regions led by a collective leader cell “front.” Finally, we demonstrate label-free imaging of millimeter-scale honeycomb-shaped tissues without the multi-image stitching often required for conventional microscopy. We utilize mutual capacitance measurements with electrically-programmable electrode spacing to reconstruct topographical features of these engineered tissues. Overall, CMOS capacitance imaging arrays enables label-free imaging spanning from single cells to large tissues, in a portable and scalable format for settings where optical microscopy may be difficult to access.

## I. INTRODUCTION

Dynamic measurements of animal cell morphology, migration, and proliferation are crucial for understanding tissue homeostasis, wound healing, and disease progression [1]. In practice, cells are tracked from a series of time-lapse images by first segmenting the cell positions (and shapes) and then linking these cell positions across time into trajectories [2]. However, accurate reconstruction of cellular motion from optical imaging is confounded by changes in cell shape, cell division or death, or cells entering or leaving the field of view. Moreover, adherent cells are thin and optically transparent, which can frustrate reliable cell segmentation and tracking using bright-field imaging, phase contrast imaging, and other label-free microscopy techniques [3]. Fluorescent labeling of animal cells permits more uniform brightness and easier segmentation, but frequent time-lapse imaging can result in photobleaching and phototoxicity [4]. Finally, there is a mismatch in length scale between single cell migration and proliferation (∼10 *µ*m) and tissue-scale reorganization (100 *µ*m to mm), which often greatly exceed a single optical field of view. Thus, imaging tissue-scale reorganization over large regions often requires sequential imaging of partially overlapping regions using a motorized stage, followed by digital stitching into one combined image.

Electrical measurements based on impedance or capacitance sensing offer one possible route to label-free, portable, low-power cell measurement platforms [5]. Early pioneering work measured how animal cells adhered [6] and migrated along planar gold electrodes using electric cell-substrate impedance sensing (ECIS). Many early demonstrations of this technology used electrodes (∼250 *µ*m) that were much larger than single cells (∼10 *µ*m), producing only aggregate measurements from a population of cells. Although ECIS is now an established commercial technology (e.g. Applied Biophysics [7], Agilent xCELLigence [8]), many of its implementations are focused on population-level readouts and cannot resolve single-cell behaviors. Modern semiconductor technology offers opportunities to scale up electrical cell culture monitoring with extremely dense microelectrode arrays (MEAs), integrated with complementary metal-oxide-semiconductor (CMOS) circuits. By combining microelectrodes with active electronic addressing and readout, CMOS MEAs offer high spatio-temporal resolution, high-throughput measurement, and additional opportunities for stimulation and manipulation of cellular function. Key demonstrations of such chips with thousands of electrodes have focused on neural recordings [9–12], but the technology can be broadly applied to many other cell types as well [13–19]. Capacitance is a particularly promising readout, as it is inherently sensitive to samples on the surface; adherent cells obstruct the electrode, acting as insulators enveloped by a polarized layer of counterions from the aqueous solution [20].

Individual animal cells use cell-surface adhesion to drive shape changes and directed migration (on planar surfaces) [21], which warrant surface-sensitive measurements in space and time. Moreover, groups of epithelial cells exhibit collective migration by using cell-cell adhesions to facilitate force transmission and larger-scale coordination [22]. For example, aggregated multicellular spheroids adhere and spread cohesively on substrates as a competition of cell-cell and cell-matrix adhesion [23–26]. Further, individual cells can detach and disseminate from this collective front via an epithelial-mesenchymal transition (EMT), associated with a loss of cell-cell attachment and strengthened cell-matrix adhesion [27]. Cells exhibit pronounced differences in shape, migration speed, and directional persistence when transitioning between epithelial and mesenchymal states [28–31].

Here, we show label-free single cell tracking using a large-scale CMOS capacitive imaging array with *>*1 million pixels and *>*1 cm^2^ active area. We demonstrate that individual mesenchymal cells can be segmented based on capacitance, so that their motility and proliferation can be extracted using optical flow. We then analyzed the adhesion and spreading of multicellular spheroids of epithelial cells, profiling how the collective front transitions to individual dissemination. We further demonstrate high resolution imaging of millimeter sized honeycomb microtissues without image stitching, using mutual capacitance to visualize topographical features. Overall, we expect CMOS capacitance imaging platforms will be widely applicable for label-free tracking of adherent animal cells, contributing new insights into collective cell migration in development and disease in settings where optical microscopes may not be easily accessible.

## II. METHODS AND MATERIALS

### A. CMOS-MEA device fabrication and operation

The capacitance sensor is a custom integrated circuit fabricated using a commercial 180 nm CMOS process. The sensor has an array of 1024 *×* 1024 pixels on a 10 *µ*m grid, which are individually addressable through row/column control signals. Capacitance measurements are made by switching the electrode voltage at MHz frequencies using a network of transistors within each pixel, similar to those in earlier generations of custom impedance imaging sensor arrays [32–34]. Spatially-resolved capacitance images are made by rapidly scanning the address across the array to sequentially measure each pixel. Acquiring a full frame 1024 × 1024 capacitance image takes less than 1 minute. The sensor chip was wirebonded to a small printed circuit board module and connected to a custom data acquisition board. The chip is controlled through an FPGA, which connects via USB to a Python environment (see Note S1 for further details). We further mounted a plastic ring around the chip surface to contain aqueous media for the adherent cells. After cell seeding, the chip was placed within a second plastic container to maintain aseptic conditions and transferred into a cell culture incubator with 5%CO_2_, 37°C, and high humidity.

### B. Cell Culture and Seeding on the CMOS-MEA device

Human mammary epithelial cells (MCF-10A) and the triple-negative breast cancer (TNBC) cell line (MDA-MB-231), transfected with the Z-CAD dual sensor expressing d2GFP and dRFP, were a generous gift from M.J. Toneff and J.M. Rosen. The Z-CAD dual sensor consists of a destabilized GFP linked to the ZEB1 3’ UTR and an RFP driven by the E-cadherin (CDH1) promoter. In addition, MDA-MB-231 cells expressing cytoplasmic GFP were a gift from Daniel Haber (Massachusetts General Hospital).

MCF-10A cells were maintained in growth medium composed of DMEM/F12 (Invitrogen, 11965-118) supplemented with 5% horse serum (Invitrogen, 16050-122), 20 ng/mL human epidermal growth factor (R&D Systems, 236-EG), 10 *µ*g/mL insulin (Sigma Aldrich, I-1882), 0.5 µg/ml hydrocortisone (Sigma Aldrich, H-0888), 100 ng/mL cholera toxin (Sigma Aldrich, C-8052), and 1% penicillin-streptomycin (Invitrogen, 15070-063). MDA-MB-231 cells were cultured in DMEM (Corning, 10-013-CV) supplemented with 10% heat-inactivated fetal bovine serum (Cytiva, SH30071.03) and 1% penicillin-streptomycin (Invitrogen, 15070-063).

Prior to cell seeding, the CMOS-MEA surface was first covered with 100% ethanol to prevent bubble formation in the shallow channels. The ethanol was then thoroughly rinsed off with deionized (DI) water at least three times to ensure complete removal. The surface of the chip was then covered with phosphate buffer saline (PBS) (Cytiva, SH30256.01) prior to cell seeding. Cells were seeded at two target densities, depending on the experimental condition: low density (5 × 10^4^ / cm^2^) and high density (5 × 10^5^ / cm^2^). To seed the cells, the PBS was gently aspirated off without completely drying the surface, followed by the addition of cells in culture medium.

### C. Live Imaging using Fluorescence Microscopy

For comparisons with standard tissue culture dishes, we used a Nikon Eclipse Ti fluorescence microscope equipped with a motorized stage, direct illumination LED (Thorburn), a multi-channel light source (Lumencor Spectra-X), spinning disk confocal unit (CrestOptics X-Light V2), CMOS camera (Andor Neo), and a 10*×* Plan Apo opjective (NA 0.75). Cells were maintained in a microscope-mounted incubator (In Vivo Scientific) under controlled conditions of 37°C, 5% CO2, and humidification. NIS Elements software was used for automated image acquisition. Image post processing was conducted using ImageJ.

### D. Cell segmentation and tracking from capacitance images

Adherent cells were segmented using a fixed intensity threshold on the capacitance image. For example, a threshold value of 63 fF was applied to the single-cell experiment results. After thresholding, segmentation artifacts were removed by particle-size filtering, excluding connected components smaller than 2 pixels. Two mask sets were generated for separate downstream analyses: non-dilated masks were used to quantify cell spreading area (Supplementary Fig.S7), whereas dilated masks were used for tracking to reduce boundary fragmentation that could otherwise be assigned as new trajectories (Fig. 3). For the tracking masks, dilation was followed by hole filling and watershed segmentation to merge disconnected boundaries while preserving separable cell regions. All mask-generation steps were performed in ImageJ.

Segmented binary mask stacks were used to extract cell centroids, enabling the reconstruction of single-particle trajectories. To preserve the local spatial context of cells, a *k*-nearest-neighbor graph was constructed for each frame using centroid-to-centroid distances. For each cell, up to five neighboring cells were included, with neighbors limited to a maximum distance of 30 pixels, corresponding to 300 *µ*m. Because the images contained crowded cells with dynamic changes in shape and position, direct nearest-neighbor matching between masks in consecutive frames was not sufficient for accurate tracking. Therefore, optical-flow-based position prediction was used to guide the matching between previous and current cell positions.

To predict cell displacement between consecutive frames, dense optical flow was computed using the Farnebäck algorithm (OpenCV) with the following parameters (*pyr_scale_* = 0.7, levels = 0.5, window size = 11, iteration = 3, *poly_n_* = 5, *poly_sigma_* = 1.2). For each active trajectory, the next centroid position was predicted by combining two terms: an optical-flow-based displacement term and a neighborhood-graph-based prediction term derived from the relative positions of neighboring tracks. The final predicted position was calculated as a weighted average, with a weight of 0.9 assigned to the optical-flow prediction and 0.1 assigned to the neighborhood-graph prediction. Predicted track positions were then matched to observed mask centroids in the next frame using a distance-based cost metric and the Hungarian assignment algorithm. Unmatched tracks were not immediately removed; instead, they were propagated as gap-filled trajectories for up to 8 frames, corresponding to approximately 160 minutes, and were terminated if they remained unmatched beyond this maximum gap length. Unmatched observed masks were initialized as new tracks, which may correspond to newly appearing cells, separated cells, or mask regions that became detectable after transient fragmentation. Potential merge and split events were recorded when multiple predicted tracks were close to a single observed mask, or when a single predicted track was close to multiple observed masks, respectively. Final outputs included centroid trajectories, event-candidate records, frame-wise observations, and neighbor-distance time series.

### E. Fluorescence microscopy of fixed cells for comparison with capacitance image

Following capacitance imaging of motile cells, the cell-seeded CMOS sensor was removed from the incubator and fixed with 4% paraformaldehyde (Fisher Scientific) in PBS for 20 minutes at room temperature. The fixed cells were then washed and stored in PBS at 4°C prior to imaging. Before image acquisition, the fluid chamber well surrounding the CMOS sensor was removed and replaced with a coverslip, ensuring a thin layer of PBS remained between the sensor and the coverslip. Fluorescence imaging was performed using a Nikon SMZ18 stereo microscope equipped with a 1.6× objective lens, a dual-band GFP/mCherry filter, and a DS-10 camera. The CMOS sensor was mounted on an automated XY stage controlled by a Mad City Labs microcontroller. To map the coordinates of the corresponding capacitance images (acquired in self-capacitance mode with a neighbors = 4 configuration), fluorescence images were captured systematically along the sensor. The four corners of the CMOS sensor served as fiducial markers to guide automated, fixed-distance translations along the sensor boundary.

### F. Spheroid and Honeycomb Tissue Preparation

Sub-confluent monolayers of MDA-MB-231 were trypsinized with 0.05% Trypsin-EDTA (HyClone ™, SH30042.02) then resuspended in culture medium to reach a concentration of at least 5.0 *×* 10^6^ cells/ml. Cell suspensions were then added to non-adhesive agarose micromolds (Microtissues, SKU: 24-35) drop-by-drop. The final cell seeding density was aimed at ∼7,200 cells/spheroid, or 250,000 cells/micromold. To form MDA-MB-231 cell spheroids with stronger cell–cell junctions, cells were pretreated with doxycycline (2*µ*g/mL; Sigma-Aldrich, D9891-1G) to drive E-cadherin expression for 2 days before trypsinization. The doxycycline treatment was continued during the formation of the spheroid. After seeding, cells were allowed to settle into the agarose micromolds for 30 minutes. The agarose microwells were then filled with supplemented culture medium and centrifuged at 1200 rpm for 3 minutes. Microwells were transferred to an incubator for 4 days so that cells could aggregate into spheroids. Spheroids were then harvested from the agarose micromolds by gentle pipetting and transferred to the CMOS-MEA device.

For honeycomb tissue formation, sub-confluent MCF-10A monolayers were dissociated using 0.05% Trypsin-EDTA (HyClone™, SH30042.02) and Accumax (Sigma-Aldrich, AM105), and then resuspended in MCF-10A culture medium. The cell suspension was seeded into a non-adhesive agarose micromold (Microtissues, SKU: 24-H) at a target density of 140,000 cells per honeycomb. The chambers were filled with culture medium and incubated for 2 days to allow tissue compaction. Honeycomb tissues were then harvested from the agarose micromolds by gentle pipetting and transferred to the CMOS-MEA device.

### G. COMSOL Simulation of Spheroid Capacitance

The capacitance simulations were performed in COMSOL Multiphysics using the Electrostatics Module. The model consisted of three main components:

- *2D array of electrodes.* The terminals were sized 8 *µ*m *×* 8 *µ*m and spaced 2 *µ*m apart, representing one row of electrodes in the capacitance-sensing pixel array.
- *Sample geometry.* For each of the models (Fig. 5D, Fig. S14B, and Fig. S12), an idealized sample geometry was created with rectangular and spherical components. Each test structure was positioned directly above the terminal array.
- *Surrounding medium.* The terminal array and test geometries were positioned at the bottom of a large block that represents the surrounding liquid medium (2000 *µ*m *×* 2000 *µ*m *×* 1000 *µ*m). The dimensions of this block were chosen to be much larger than those of the test structures to reduce boundary and edge effects.

The sample geometries in both Fig. 5D and Fig. S14B were assigned a dielectric constant *ϵ_r_* = 20 to approximate the cell spheroids and tissues. In addition, thin high-dielectric layers (*ϵ_r_* = 100, shown in purple) were included to model charge accumulation at the interface. The terminal array and test structures were immersed in water (*ϵ_r_* = 75).

A zero-charge boundary condition was applied to all outer boundaries of the surrounding medium. The initial electric potential (phi) was set to zero for all simulation domains, and charge conservation in fluids was applied to all domains. Different virtual pixel sizes were implemented by grouping neighboring terminals into larger terminal groups.

A stationary source sweep was used in the study to sequentially excite each terminal group. By biasing the terminal groups one at a time, the Maxwell capacitance matrix was extracted, providing the mutual capacitance between all virtual pixel groups. Finally, the mutual capacitance measured directly under the test structure (C_m_), was compared with a measurement far from the test structures, where only the surrounding medium was present (C_m,media_). The normalized capacitance was then calculated as the ratio

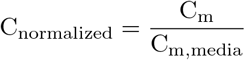

## III. RESULTS

### A. Device implementation of a one megapixel CMOS-MEA capacitance imaging sensor

We designed a custom one-megapixel CMOS-MEA sensor chip which hosts an array of 1024 *×* 1024 electrode pixels at a 10 *µ*m grid pitch, creating an active sensing area of 10.24 mm *×* 10.24 mm = 104 mm^2^ (Fig. 1A). This sensor is a new chip which adds more pixels, improved calibration features, new measurement modes, and higher resolution than our previous designs [32–35] (see Note S1 and Fig. S1-S6 for further details). This design enables large area measurements with cellular-scale resolution, ranging from spheroid spreading at the scale of hundreds of pixels (∼1000 *µ*m) down to single adherent cell morphology with multiple pixels (∼ 30 *µ*m) (Fig. 1B,C). The custom sensor chip is fabricated in 180 nm CMOS technology with overall dimensions of 12.5 mm *×* 12.6 mm. The chip is wire-bonded to a small printed circuit board, and the bondwires are encapsulated with epoxy. We typically attach a small open fluid chamber around the sensor using silicone elastomer (Fig. S1A). The top aluminum metallization layer is chemically removed from the pixels, exposing the titanium nitride layer underneath as the sensing electrode surface [32]. The surface of the CMOS sensor is in direct contact with aqueous media and the adherent cells.

**FIG. 1.**
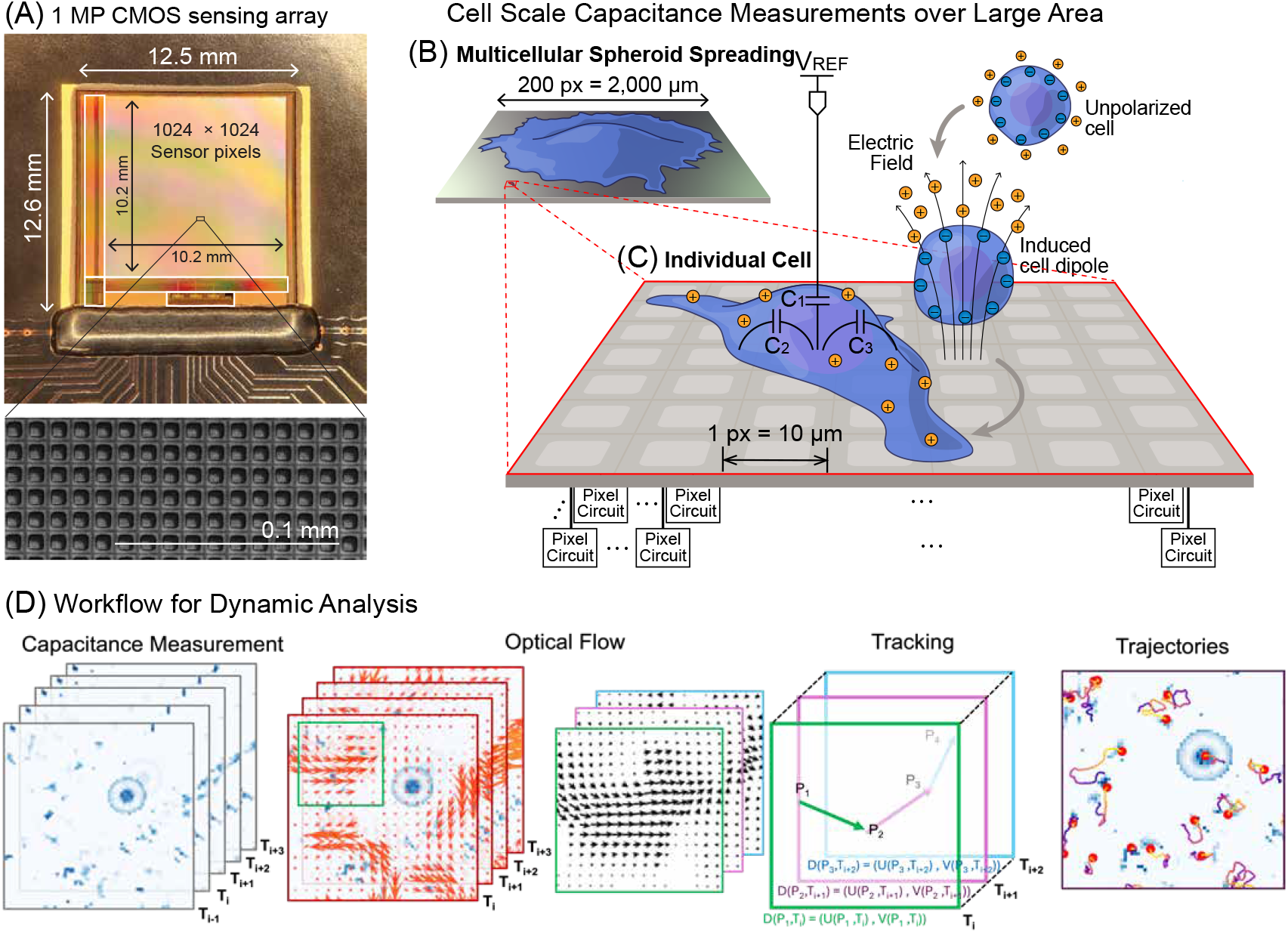
Scheme for capacitance imaging using CMOS-MEA sensor. (A) Representative optical image of the custom 1 megapixel CMOS MEA sensor (top) and an electron microscope image of the 10 *µ*m electrode pixels (bottom) (B) A multicellular spheroid spreading across the CMOS MEA surface can occupy thousands of electrodes, spanning several millimeters. (C) Individual cells typically occupy a few electrodes on the order of tens of microns. The chip measures local changes in capacitance which correspond to changes in ion redistribution around cells in response to radio-frequency pulsed electric fields. (D) Velocity fields of the cells are extracted from the capacitance measurement data using optical flow. Cell trajectories are then reconstructed by updating the position of the capacitance map of adhesive cells with velocity fields across consecutive time points.

This CMOS-MEA sensor permits different capacitance sensing modes to measure adherent cells, which act as insulating structures that block the electrode surface, covered by an electrical double layer of counterions (Fig. 1B,C). In the simplest ‘self-capacitance’ mode, one pixel is repeatedly charged and discharged while the sensor measures the average charge transfer required to reach the specified voltage (see Supplementary Information and Fig. S2). The self-capacitance is the sum of the vertical coupling to the bulk, plus the lateral coupling to other nearby pixels [34]. The sensor also supports a modified ‘shielding’ self-capacitance mode in which the neighboring pixels are switched in phase with the sensing pixel, thereby shielding the lateral fringe fields so that only the vertical fields remain. Since some of the lateral coupling occurs within the sensor and very close to the surface, the shielding mode tends to improve the capacitive contrast of materials above the sensor and moderately enhance the signal to noise ratio (Fig. S5). Finally, the sensor can perform mutual-capacitance measurements between electrically-programmable rectangular *sets* of electrodes[36]. For example, we can measure the mutual capacitance between two different 2*×*2 pixel groups, or two different 3*×*3 groups. While the microelectrodes are physically on a 10 *µ*m pitch, these measurements can approximate the mutual capacitance between larger ‘virtual pixels’, for example 20 *µ*m (*V* = 2) or 50 *µ*m (*V* = 5). In each of these sensing modes, the measurements are performed serially as the addresses raster through the array. We usually acquire 24,000 pixels/sec, translating to less than 1 minute per frame, which is sufficient for time-lapse cell culture imaging. The charge/discharge cycles operate at MHz frequencies, often yielding lower absolute capacitance compared to more traditional kHz impedance measurements, but MHz frequencies can also improve the detection depth and material contrast due to reduced counterion screening [15].

Our workflow for dynamic analysis of cell migration was based on sequential capacitance measurements (e.g. every 15 min), analogous to time-lapse imaging in optical microscopy (Fig. 1D). For each capacitance “snapshot”, adherent cells exhibited lower capacitance than the surrounding aqueous media, resulting in a strong signal relative to background. An optical flow algorithm was then applied to sequential capacitance images in order to reconstruct motion fields [37]. From these pixel-wise velocity vectors at each time frame, we then reconstructed single-cell trajectories by propagating from initial cell positions over time.

### B. Single cell detection using capacitance imaging

We compared capacitance and fluorescence imaging for single cell detection and the extraction of shape features, noting that the 10 *µ*m CMOS-MEA sensor pixels are smaller than a typical adherent animal cell. Using cultured breast cancer cells (MDA-MB-231) that exhibit mesenchymal migration, we observed that cells exhibited lower capacitance than the surrounding unoccupied regions (Fig 2A). We then immediately fixed these cells (which also express green fluorescent protein) and performed optical imaging using an upright optical fluorescence microscope (Fig 2B). Our comparison of capacitance and fluorescnece imaging showed good qualitative agreement for features such as the cell nucleus (Fig 2C,D). However, peripheral subcellular features such as the lamellipodium exhibited higher contrast in capacitance imaging than in fluorescence, perhaps since these parts of the cell obstruct the electrode surface but are relatively thin (Fig 2C,D). Indeed, the intensity profile for capacitance imaging shows a sharper threshold for segmenting foreground features relative to background features (∼ 60 fF, Fig 2E), whereas the fluorescence threshold was less pronounced (∼50 (a.u.), Fig 2F). For further comparison between the capacitance and fluorescence images, we matched the spatial resolution by upsampling the capacitance image by 10-fold using bilinear interpolation. We then trained a deep learning cell segmentation algorithm (CellPose cyto3 [38]) using ∼1,000 manually annotated cell masks, which was applied to segment both capacitance and fluorescence images (Fig 2A,B). We observed reasonable agreement in the approximate position of cells segmented in capacitance and fluorescence images. However, the boundary detection in fluorescence images appeared to systematically underestimate cell size relative to capacitance images (Fig 2G), likely due to poorer segmentation of peripheral features (Fig 2A,B). In comparison, shape metrics such as eccentricity were more correlated between capacitance and fluorescence imaging, without any systematic bias (Fig 2H). We note that the segmentation of fluorescence cells on a CMOS MEA chip is degraded by autofluorescence from the sensor surface materials and grid texture. As a result of this artifact, the segmented cell shape based on fluorescence was scaled down slightly relative to the corresponding cell shape based on capacitance, resulting in decreased area but comparable eccentricity. Principal component analysis (PCA) further confirmed that capacitance and fluorescence derived cell masks exhibited comparable shape features, with substantial overlap in the morphological feature space (Fig. S7A,B,C,D). We further confirmed that cell boundaries could be segmented from the raw capacitance images without upsampling the spatial resolution (Fig. S7E), since the high signal to noise and uniform contrast across the field of view permitted simple global thresholding (∼ 61 fF). Notably, the distribution of major axis lengths of MDA-MB-231 cells was comparable for capacitance images (with or without upscaling) relative to phase images acquired of these same cells adherent to tissue culture plastic (Fig. S7E,F). In comparison, the major axis lengths from fluorescence imaging (on the CMOS-MEA) was skewed towards smaller values, likely due to poorer boundary segmentation.

**FIG. 2.**
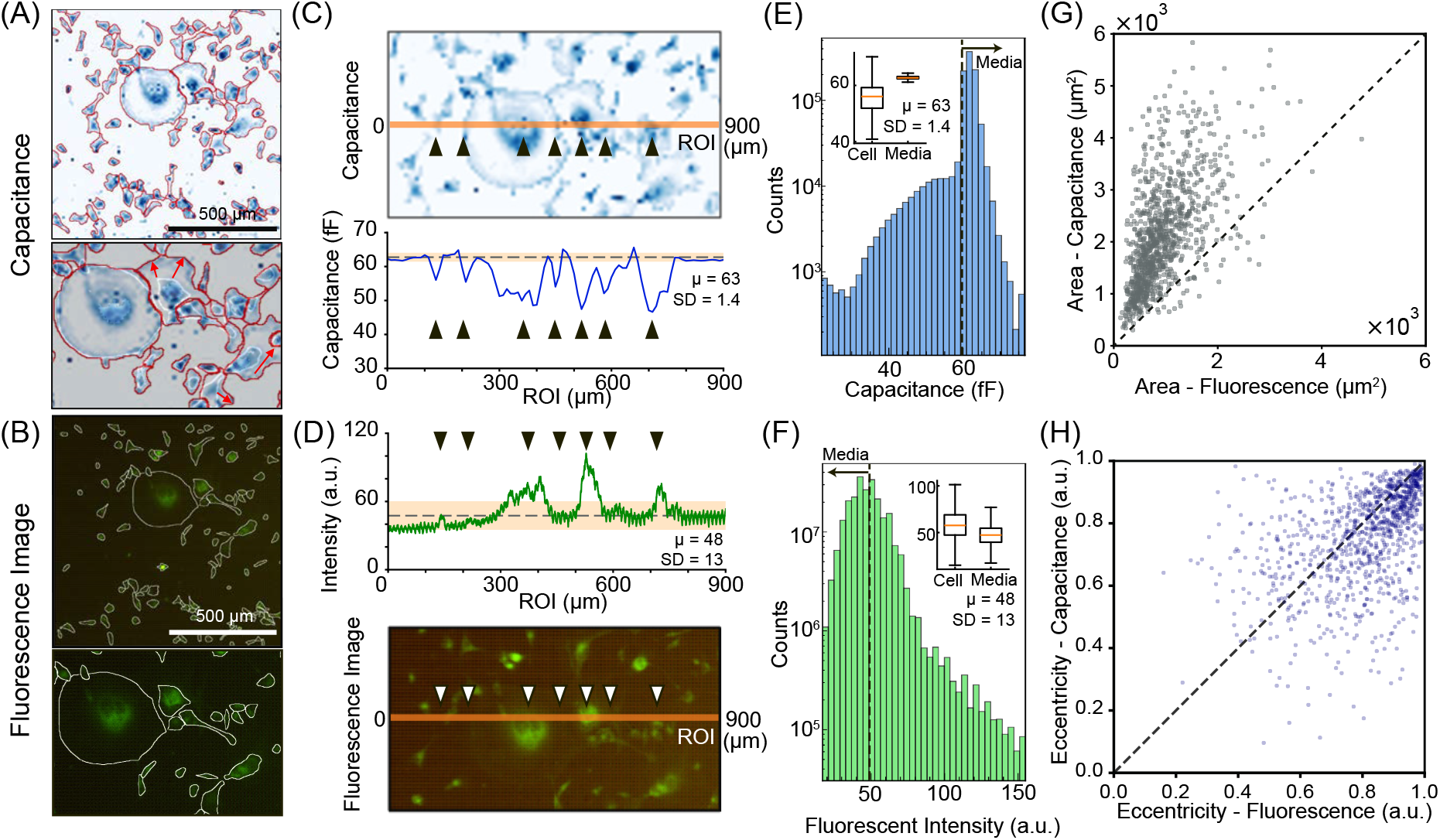
Comparison of cell segmentation using capacitance and fluorescence imaging. (A, B) Capacitance and fluorescence images of GFP-expressing MDA-MB-231 breast cancer cells. Boundary outlines were extracted from both images using deep-learning cell segmentation (capacitance, red; fluorescence, white). Arrows indicate discrepancies between the boundaries identified from optical images and those obtained from capacitance measurements. (C, D) Matched regions of interest and linear intensity profiles for both capacitance measurement (C) and fluorescence intensity(D). Arrows denote subcellular features of interest, including the cell boundary interface, nucleus, and cell body. The dotted line indicates the mean value in the media region, and the orange box represents the standard deviation of the capacitance in the media region. (E,F) Histogram of capacitance values and fluorescence intensity within the entire region. Inset shows the distributions of capacitance, and fluorescence values for the cell interior, defined by the cell mask, and for the surrounding media region. (G, H) Comparison of cell area (G) and eccentricity (H) between CMOS chip and optical images. Each dot represents a matched pair of cell masks obtained from capacitance and fluorescence images. The dotted lines indicate *y* = *x*, as a reference line.

**FIG. 3.**
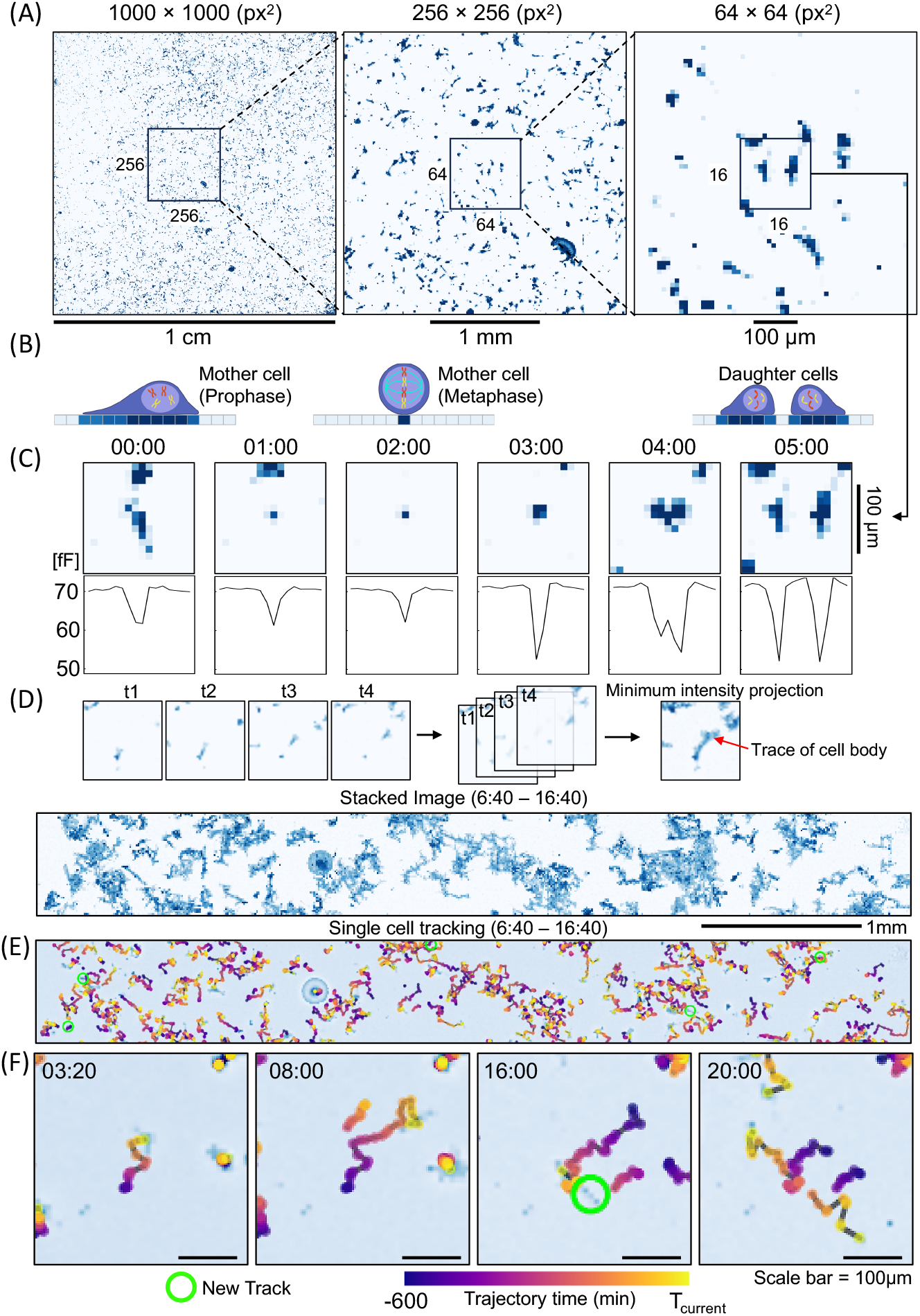
Large field-of-view cell tracking using time-lapse capacitance imaging. (A) Large field-of-view (FOV) allows continuous segmentation and tracking of single cell dynamics, capturing rare events such as cell divisions and rapidly migrating cells. The smallest ROI window(16 × 16px^2^) marks a cell division event. (B) Illustration of the cell division process and corresponding changes in the capacitance map. (C) Changes in the capacitance map and line profile of ROI center during cell division. (D) Minimum intensity projection of capacitance images over 30 frames (10 h). (E) Single-cell trajectories tracked within the same ROI as in (D), overlaid on the capacitance image at 16:40. Bright colors indicate current positions, whereas darker colors indicate previous positions. (F) Representative single-cell trajectory showing cell division at 16:00, with daughter-cell tracks emerging after division. The green circle indicates the initiation of a new track.

We further verified the capacitance image quality with a second cell type, a non-transformed mammary epithelial cell line (MCF-10A) (Fig. S8). These cells were initially plated at a sparse density of ∼ 20%, which proliferated into a confluent monolayer over 43 h. The proliferation rate of these MCF-10A cells was consistent with typical values on tissue culture plastic (doubling time ∼20 hours), and confirmed the biocompatibility of cells with the CMOS-MEA chip [39].

### C. Tracking individual cell migration from capacitance imaging

Next, we investigated whether cell trajectories could be reconstructed from time-lapse capacitance imaging. As a representative example, the full frame capacitance snapshot is successively “zoomed in” from 1024 × 1024 pixels (1 cm) down to 256 × 256 pixels (2.56 mm) and then 64 × 64 pixels (640 *µ*m) (Fig. 3A). We again plated breast cancer cells (MDA-MB-231) at a sparse density of ∼5%, and performed capacitance imaging for the next 20 h. Notably, we could identify thousands of cells dispersed across the CMOS-MEA sensor surface, each typically spanning multiple pixels.

Strikingly, cells remained visible in the capacitance imaging even during division events. For example, an initially spread “mother” cell gradually rounded into a compact morphology over 2 h (i.e. prophase to metaphase) (Fig. 3B,C). The cell then spread again and eventually divided into two “daughter” cells, which was apparent from a linescan where a single valley at 3 h progressively resolved into 2 distinct valley by 4 h (Fig. 3B,C). Such cell division events are very challenging to resolve using fluorescence imaging due to the dramatic shape changes and transient loss of fluorescence marker expression, resulting in the cell temporarily “disappearing.” [40]. These proliferation events are relatively infrequent and unpredictable, but are more easily captured using label-free capacitance imaging (without photodamage) with many cells in a large area.

The stability of the capacitance signals for adherent cells facilitated single cell tracking, which we implemented by combining optical flow with cell segmentation in each frame. Briefly, the position of each cell in the next frame was predicted using the optical-flow displacement vector at the mask centroid, and the predicted positions were assigned to segmented masks by Hungarian matching (see Methods for further details). When capacitance measurements were read out at sufficiently fast time intervals (relative to cell motion), these tracking steps could be integrated over time to reconstruct cell migration trajectories. To validate the reconstructed trajectories, we compared them with sequentially overlaid capacitance images (Fig. 3D,E, Fig. S9). The trajectories were in good agreement with the cell traces observed in the “stacked” capacitance images, confirming accurate single-cell tracking. This tracking capability was also evident in the reconstruction of newly generated trajectories, including daughter-cell trajectories after division, enabled by accurate mask segmentation and high-resolution migration tracking (Fig. 3F).

We applied the same optical flow analysis to phase contrast microscopy images of MDA-MB-231 cells adherent to tissue culture plastic in a multiwell plate (Fig. S10A, B). We measured comparable migration metrics (e.g. average cell speed, directionality, and diffusion coefficients) for cells on the tissue culture plastic relative to the CMOS-MEA sensor. Over 24 h, we further estimated that 5-10% of the cells imaged using phase contrast microscopy would migrate out of the 1600 *×* 1300 *µ*m^2^ field of view (Fig. S10D,E,F,G). This issue of cells leaving (and entering) the field would likely be further exacerbated for time-lapse optical imaging over longer durations. Overall, these results show that capacitance imaging enables accurate, label-free quantification of live-cell motility without tracking artifacts associated with cells disappearing due to proliferation or moving out of the field of view.

### D. Analyzing collective migration dynamics of spreading multicellular spheroids

Multicellular spheroids can be formed by aggregation of hundreds to thousands of individual cells, which can then be transferred to planar surfaces, where they disorganize and spread on planar surfaces as a competition between cell-cell and cell-surface adhesion [23, 24, 41]. Cells at the periphery of these spreading aggregates often exhibit a transition from collective to individual migration that is reminiscent of an epithelial-to-mesenchymal transition [23]. We prepared spheroids by aggregating ∼ 7,200 MDA-MB-231 cells for 4 days in low-adhesion agarose micromolds, after which they were transferred to the CMOS MEA sensor surface. These spheroids were initially ∼ 350 *µ*m in diameter, then gradually flattened and widened over the next 50 h (Fig. 4A). Based on capacitance imaging, we extracted a spreading area as a function of time *A*_spreading_(*t*) by applying a capacitance threshold (∼ 35 fF) to generate binary masks. This area was further coverted into an effective radius, determined by equivalent radius corresponding spreading area, 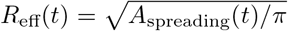 (Fig. 4B). We found that the radial expansion of the aggregate gradually slowed down over time, which could be fit to a power law, *R*_eff_(*t*) ∼ *t^α^*, where the exponent *α ≈* 0.4 (Fig. S11). This exponent is comparable to those from scaling arguments that treat “active wetting” based on the spreading of viscoelastic liquid drops. Briefly, early spreading dynamics exhibit *α ≈* 1*/*3 based on the the competition of surface energy gain and viscous dissipation (at short times), which transitions to *α ≈* 1*/*2 at later times when the liquid film can “flow” along the surface [23].

**FIG. 4.**
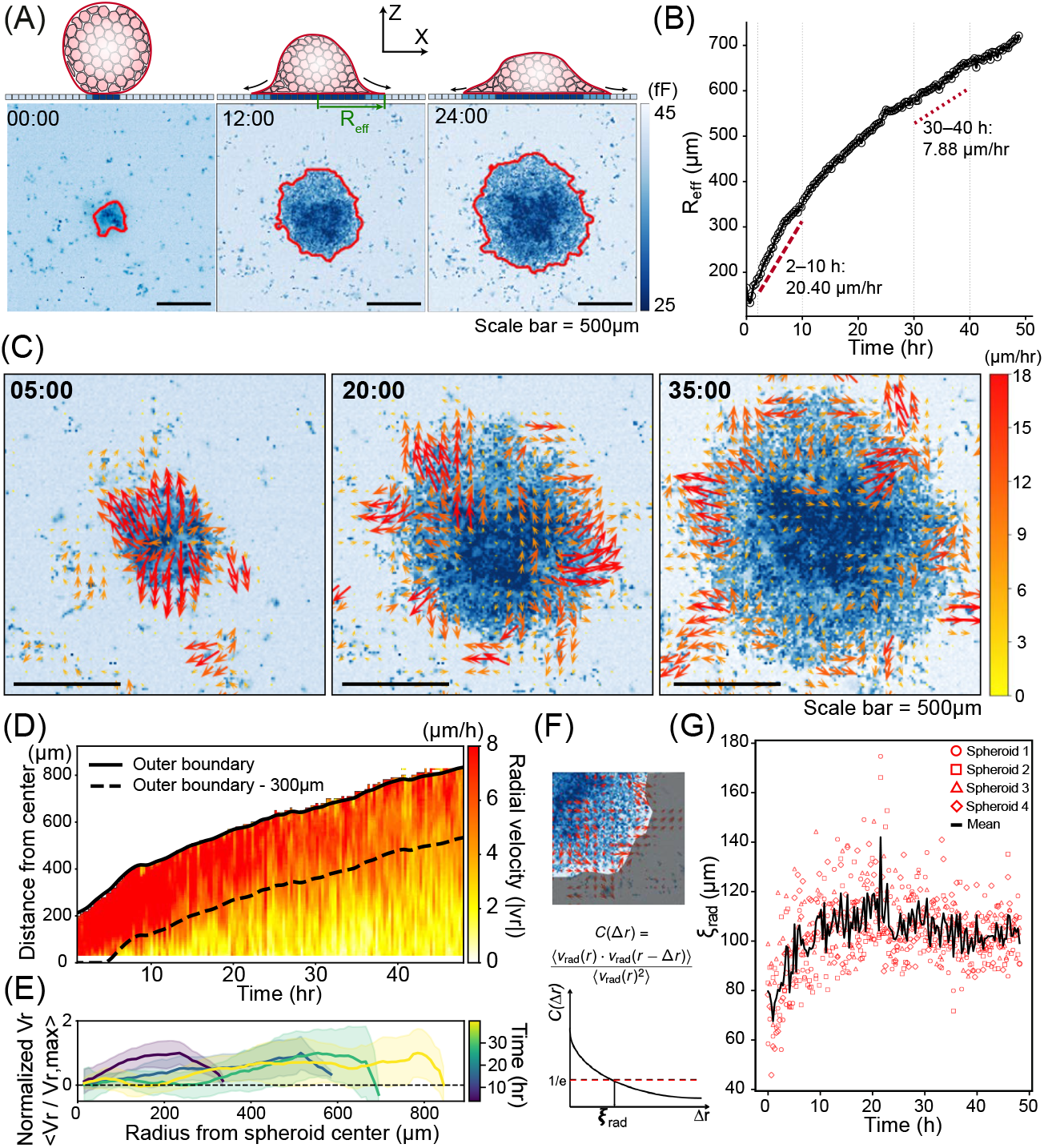
Multicellular spheroid spreading and collective migration analysis using capacitance imaging. (A) Representative capacitance images of spheroid spreading over 24 h. Red outlines indicate the spreading boundary determined by image thresholding. (B) Spreading trend of representative MDA-MB-231 spheroid over time, the effective radius(*R_eff_*) is determined by the equivalent radius corresponding spreading area. Dotted lines indicate average slopes of the lines connecting the values at 2-10 h and 30-40 h. (C) Representative images of spreading spheroids with velocity fields reconstructed using an optical flow algorithm. (D) Spatiotemporal changes in radial velocity inside of spreading domain, the solid line indicate the outer boundary of spreading, and dotted line displayed the inner 300*µ*m from the boundary. (E) Development of velocity profile from the boundary to core over time. (F) Schematic for calculating correlation length of velocity fields. (G) Correlation lengths of radial velocity inside of spreading masks.

We further used optical flow to extract the motion fields of the spreading aggregate, revealing collective cell migration was fastest at the periphery, oriented radially outwards (Fig. 4C). Nevertheless, there was also a stagnant region at the center that also increased in size over time. We visualized the radial dependence of this velocity using a kymograph, revealing this fast-moving peripheral region (“leader cells”) were typically localized to the outermost 300 *µ*m of the spreading aggregate, with a rapid dropoff in speed near the center (Fig. 4D). Indeed, the radial velocity profile appeared self-similar at different times, with a minimum at *r* = 0, increasing up to a maximum velocity near the aggregate periphery (Fig. 4E). Moreover, cells within this peripheral region gradually increased their coordination, sustaining a characteristic length scale of 100 *µ*m from 10-50 h based on the spatial correlation of the velocity field (Fig. 4F,G).

### E. Depth-Sensitive Mutual Capacitance Measurements using Virtual Pixels

The spreading of a multicellular spheroid on a surface results in a gradual decrease in aggregate height as the originally-spherical core becomes progressively flatter and a disc-shaped sheet of cells make contact and spread along the sensor surface (Fig. 4A). This represents an intriguing test case for the depth sensitivity of the chip’s different mutual capacitance sensing modes. While the physical pixels are on a 10 *µ*m pitch, we can also measure the combined mutual capacitance between two closely spaced groups of electrodes. For example, an 8 *×* 8 pixel group behaves like a larger 80 *µ*m “virtual pixel”, which we denote by *V* = 8. Since the depth of the fringing electric fields varies with the size and spacing of the electrodes [42] (Fig. 5A), different virtual pixel sizes (*V*) will have different depth sensitivity. Interestingly, at ∼ 23 h the spheroid’s single-pixel mutual capacitance (*V* = 1) was greater than 1 (normalized to the background media). However, for larger effective electrode sizes (*V ≥* 2), the spheroid capacitance was lower than the background media capacitance (Fig. 5B,C). We hypothesized that these differences occurred due to the increasingly out-of-plane fringing of the electric field. To estimate this depth sensitivity, we implemented a simple COMSOL simulation for a rectangular block with dielectric constant *ϵ* (greater than media) and varying height *H*. We sought to find the critical height above which the virtual pixel measurement is no longer able to detect the presence of a cell-like object. As the virtual pixel size increases from *V* = 1 to *V* = 8, we find that the depth sensitivity improves at the cost of lateral resolution, and that the detection depth increases from 4 *µ*m to 60 *µ*m (Fig. S12).

**FIG. 5.**
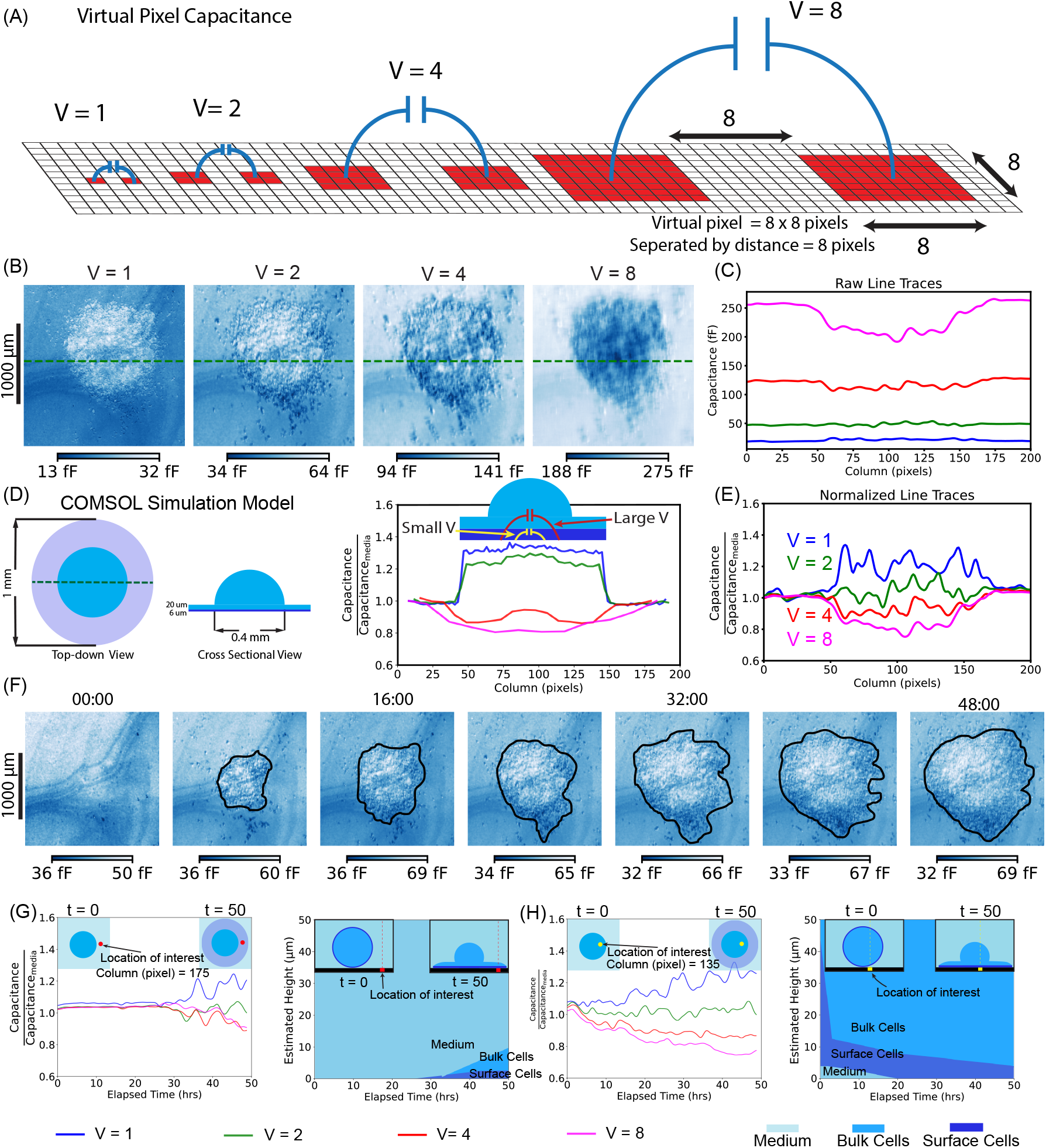
Mutual capacitance measurements using “virtual pixels” reveals topographical information about tissue height. (A) Depiction of programmable virtual pixels consisting of spatially separated sets of pixels, with varying out-of-plane electric field fringing that recovers information at varying heights. (B) Capacitance image with virtual pixel size varying 1, 2, 4, and 8 pixels at t = 23 hrs. (C) The raw capacitance line trace across the spheroid for all virtual pixel sizes. (D) COMSOL simulation for a hemispherical tissue with 200 *µ*m radius over a thinner spreading cell layer with diameter of 1 mm. (E) Line trace with normalized capacitance values plotted for virtual pixels ranging from *V* = 1, 2, 4, 8. (F) Time-lapse images of a spheroid containing MDA-MB-231 cells using virtual pixel image with size = 2. (G, H) Temporal evolution of virtual capacitance signals for a spheroid spreading at two different locations along the radial direction of the spread. Along with the estimated distribution of cell media, surface and bulk cells along the depth at different time points. All the experimental measurements of line traces were denoised via a 1D median filter (size = 3), and smoothed using a 1D Gaussian filter (*σ* = 2)

We then considered a more complex simulation geometry to better represent the experimental spheroid spreading. Briefly, we simulated the spheroid as a hemispherical cap (d = 400 *µ*m), surrounded by a planar layer (h = 20 *µ*m), over a thinner high-dielectric layer (h = 6 *µ*m) which accounts for charge accumulation near the electrode surface (Fig. 5D). Smaller virtual pixels (*V* = 1, 2) were primarily sensitive to the planar monolayer, and capacitance increased due to the surface charge [20, 43]. In contrast, the mutual capacitance for larger virtual pixels (*V* = 4, 8) is dominated by the reduced dielectric constant of the hemispherical cap, producing capacitance lower than the media background. These simulated results are in qualitative agreement with our experimental measurements (Fig. 5E).

We then considered time-lapse measurements across a range of virtual pixel sizes (*V* = 1 to *V* = 8, in Fig. 5F and Fig. S13). We focused on representative readouts from two locations, the first near the spheroid periphery (Fig. 5G) and the second associated with the spheroid center (Fig. 5H). At the peripheral location (Fig. 5G), between 24-40 h we only observe the *V* = 1 capacitance increasing, suggesting a monolayer layer of collectively migrating “leader” cells. After 40 h, we observe mutual capacitance for *V* = 4, 8 decreasing while *V* = 2 remained comparable, suggesting a thickening layer of cells. Based on the relationship with the COMSOL simulations, we can estimate the height of the cellular layers at the peripheral location as a function of time.

Closer to the spheroid center (Fig. 5H), the *V* = 1 capacitance increased after 16 h, as the cells adhered more strongly to the surface. Meanwhile, the mutual capacitance of *V* = 4, 8 gradually decreased over time, which corresponds to the spreading of the larger spheroid mass. Overall, this proof-of-concept highlights how the use of mutual capacitance measurements via virtual pixels of varying size can yield depth information, which could potentially be used for label-free imaging of three-dimensional specimens.

### F. Spatially Heterogeneous Spreading of Millimeter-Scale Honeycomb Tissues

To further validate these large area measurements, we prepared honeycomb-shaped tissues from 140,000 mammary epithelial (MCF-10A) cells. Briefly, a non-adhesive agarose microwell was replica molded to present a hexagonal array of pillars that were 600 *µ*m in diameter with 400 *µ*m separation [44], then seeded with a dispersed cell suspension. After two days, the mammary epithelial cells aggregated around each pillar to form a honeycomb shape that was 3 mm in diameter. Note that the cells essentially aggregate into interconnected toroids that surround each agarose pillar, and the cell contractility results into tissue morphologies that conform around each pillar. As a consequence, the honeycomb tissue incorporates six circular cell-free voids that correspond to the pillar locations. These honeycomb-shaped tissues were then transferred onto a transparent tissue culture plate or CMOS MEA surface for imaging (Fig. 6A). As the honeycomb tissue adhered to the surface, it gradually flattens outwards while also collapsing the circular voids (which are no longer occupied by agarose pillars) (Fig. 6B).

**FIG. 6.**
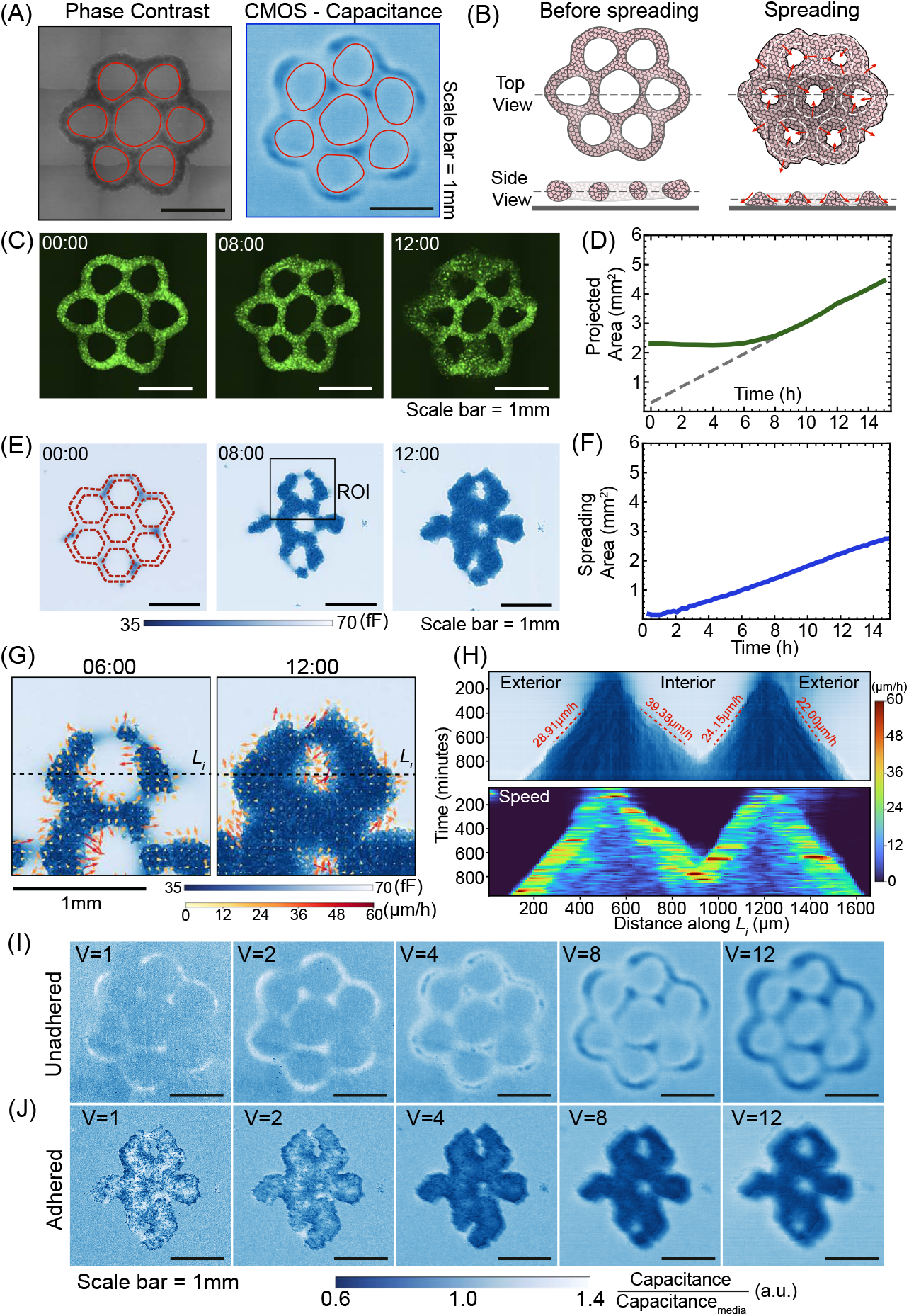
Adhesion and spreading of millimeter-sized honeycomb tissue of MCF-10A cells. (A) Comparison of honeycomb-shaped tissue morphology by phase contrast optical microscopy and CMOS mutual capacitance measurement (virtual pixel = 8), red contours indicate empty voids. (B) Schematic of the tissue spreading experiment. The dotted line indicates the central cross-section shown in the side view. (C) Spreading of green fluorescent protein expressing honeycomb tissue using epifluorescence microscopy. (D) Change in projected tissue area over time using epifluorescence imaging; the dotted line indicates the projected area of the tissue body at the initial time point. (E) Spreading of honeycomb tissue based on capacitance imaging over time; the orange dotted line indicates the approximate outline of the tissue. (F) Tissue spreading area over time measured from capacitance values. (G) Velocity field of tissue spreading within the region of interest marked in *(E)*, showing spreading dynamics in the interior and exterior regions of the tissue. Arrows indicate velocity vectors, and the colormap represents speed. (H) Kymographs of spatiotemporal changes in the tissue body, shown as the darker region, and speed along the line section (*L_i_*) marked in *(G)* with a width of 100*µ*m. Red dotted lines indicate the approximate slope on each side. (I, J) Normalized capacitance measurements obtained using different virtual-pixel sizes(V=1,2,4,8, and 12) in mutual capacitance imaging of unadhered and spreading honeycomb tissue.

We first performed fluorescence imaging of the green fluorescent protein labeled cells, revealing that the honeycomb-shaped tissue morphology remained intact after transfer (Fig. 6C). Typically, the circular voids had a diameter of ∼ 600 *µ*m separated by tissue sections with thickness ∼ 170 *µ*m. Over the next 12 h, the tissue appears to gradually spread outwards with increasing footprint. The total fluorescence intensity decreased slightly near the spreading fronts, which may indicate some thinning of the tissue with increasing area. Based on a fluorescence threshold, we further quantified the (projected) tissue area over time, which remained roughly constant at 2.5 mm^2^ for the first 8 h. Afterwards, the spreading area increased linearly at a rate of ∼ 0.3 mm^2^ / h (Fig. 6D).

In comparison, the honeycomb-shaped tissue was barely visible in the capacitance imaging at 0 h (Fig. 6E), since there was minimal cell adhesion. Nevertheless, the adhesion and spreading of the honeycomb-shaped tissue was increasingly apparent over the next 8 h. Indeed, after 12 h, the tissue had spread beyond the original tissue boundary, resembling the behavior observed in the optical images (Fig. 6E). Thus, capacitance imaging directly reveals the kinetics of tissue adhesion and spreading, starting from almost nonexistent adhesion at 0 h, then increasing linearly in time (Fig. 6F). This contrasts with the projected area extracted from fluorescence imaging, which only shows an increase once the tissue spreading on the surface exceed the largest features of the initial honeycomb shape (Fig. 6D). We note that the spreading rate was slower on the CMOS chip compared to the tissue culture dish and the CMOS chip, likely due to differences in the texture and surface chemistry. However, the linear scaling of spreading area at later times was qualitatively consistent for the tissue culture plate and CMOS-MEA.

We further examined how the honeycomb tissue spread to fill the empty interior region relative to expanding the exterior boundary. Notably, interior boundaries adjacent to empty voids are curved inward (convex), while exterior boundaries are curved outwards (concave). We again computed velocity fields using optical flow, identifying a stagnant region at the center as well as peripheral regions exhibiting coordinated migration towards unoccupied areas (Fig. 6G). Kymographs of capacitance images and speed maps along the center line (*L_i_*) of on one side of the honeycomb also revealed the spatiotemporal spreading dynamics of the tissue (Fig. 6H). Peripheral regions of coordinated motility increased in width from ∼130 *µ*m to ∼ 500 *µ*m from 4 h to 12 h. Moreover, the collective front migrating towards the interior moved faster at 39.96 *µ*m/h than the front at the exterior boundary at 28.91 *µ*m/h (Fig. 6H).

The honeycomb-shaped tissue also presents a complex topography that could be interpreted using virtual pixel mutual capacitance. For example, a honeycomb-shaped tissue that was not adhered to the surface was barely visible with *V* = 1. With increasing virtual pixel size, regions of the honeycomb that were farther from the surface could be visualized. Notably, *V* = 2 again showed a higher mutual capacitance of tissue relative to media, while *V ≥* 4 showed a lower mutual capacitance of tissue relative to media, consistent with the spheroid measurements (Fig. 6I,J). COMSOL simulations using a honeycomb-like geometry section with an offset step revealed the sensitivity of mutual capacitance to a surface layer and bulk tissue mass, respectively. In comparison, an adhered and spreading honeycomb tissue showed a lower mutual capacitance of tissue relative to media for almost all virtual pixel sizes. This scenario was treated using a COMSOL simulation of a solid mass of cells adherent to the surface, resulting in similar trends for mutual capacitance across different virtual pixel size and time-lapse measurements (Fig. S14A,B). Overall, these results highlight the capabilities of this CMOS-MEA sensor for capturing complex millimeter-scale tissue dynamics without labeling or additional imaging components, while resolving curvature-dependent spreading and regional differences in collective migration.

## IV. DISCUSSION AND CONCLUSION

We demonstrate a CMOS-MEA capacitive sensor array for label-free electrical tracking of live cells across a *>* 1 cm^2^ area over several days. As adherent cells obstruct the electrode pixels, we measure a pronounced change in capacitance with good signal to noise ratios. We further show that these 10 *µ*m wide pixels are sufficient to resolve subcellular features of single adherent cells, particularly to define the nucleus and boundary. Although the results presented here focus on epithelial and mesenchymal cells, this label-free capacitance imaging platform is cell-agnostic. We segment single cells in these capacitance images using established deep learning algorithms (e.g. CellPose [38]), yielding cell area and length measurements that are consistent with those measured for the same cell line using optical microscopy (on tissue culture plastic). We note that capacitance imaging using 10 *µ*m pixels has coarser spatial resolution than a typical optical microscope, making the resulting images appear more “pixelated.” An exciting prospect for future work is to upscale capacitance images based on deep-learning super-resolution algorithms trained on fluorescence microscopy images [33, 35, 45]. Moreover, the 10 *µ*m pixel pitch is not an intrinsic limitation, and we anticipate that reducing the electrode pitch to ≤ 5 *µ*m should be possible in other mature CMOS technology nodes (i.e. 90/65 nm). This may be useful for applications with immune cells such as lymphocytes or neutrophils, which are smaller than epithelial cells and weakly adherent to surfaces.

We further show that single cell tracking is easily implemented using optical flow algorithms, since the cell segmentation of capacitance images is robust. Time-lapse imaging every 15 minutes was sufficient to recover cell trajectories that are consistent with the same cell line measured on tissue culture plastic. Moreover, we can continuously resolve single cells even during division events, which is often problematic for fluorescence imaging. While faster frame rates were not necessary for these experiments, another intriguing possibility for future work is to implement adaptive imaging based on real-time cell segmentation and tracking [45]. For example, pixels can be selectively read out in close proximity to regions of low capacitance (i.e. adherent cells), while unoccupied regions are ignored. Our preliminary results further demonstrate additional topographical information can be recovered using mutual capacitance with “virtual pixels.” Such capacitance imaging with increased distance from the surface is promising, although there is a trade-off with the loss of lateral resolution as well as slower readout of multiple virtual pixel sizes compared to only capacitance of single pixels. We also anticipate that deep learning may be useful to integrate the readouts of self-capacitance and mutual capacitance to reconstruct three-dimensional tissue architecture with higher spatial resolution.

Although the present work focused on cell adhesion to planar substrates, we envision that mutual capacitance imaging could also be used for organoids or organ-on-chip platforms where cells are fully embedded in a compliant three-dimensional hydrogel [46]. Multicellular spheroids also exhibit complex collective and individual invasion behaviors in 3D hydrogel matrix [47, 48], which we anticipate can be resolved label-free using capacitance imaging since these hydrogels have very high water content. It may also be useful to scale up these CMOS-MEA sensors for a multiwell plate platform, as recently demonstrated by Abbott and coworkers [49], both for experimental convenience and greater compatibility with laboratory automation. Alternatively, these sensors may be useful for point-of-care diagnostics where optical microscopes are difficult to access, and opportunities would exist for integration with microfluidic sample processing [50].

In conclusion, we have demonstrated label-free imaging of adherent animal cells using a CMOS-MEA capacitance sensor with 1 million pixels at 10 *µ*m pitch. As a representative case study, we demonstrated segmentation and tracking of breast cancer cells exhibiting mesenchymal migration. We further investigated the spreading and disorganization of multicellular spheroids of epithelial cells, revealing a fast moving peripheral region of “leader cells” surrounding a more stagnant interior. We demonstrated that measuring mutual capacitance between programmable sets of pixels can yield thickness information about three-dimensional spheroids, and we further applied this technology to characterize large honeycomb shaped tissues that were several millimeters in diameter. We anticipate that non-optical capacitance imaging will enable new insights into collective and individual cell dynamics with biomedical relevance for inflammation, wound healing and tumor progression.

## ACKNOWLEDGMENTS

We acknowledge funding from Brown University for a Division of Research Seed Award (JKR, IYW), a School of Engineering Hazeltine Innovation Award (JKR) and a Hibbitt Postdoctoral Fellowship (JK). This work was also supported in part by the National Science Foundation under Grant 2027108 (JKR), as well as the U.S. National Institutes of Health R01GM140108 (HT, JK, AV, IYW).

## AUTHOR CONTRIBUTIONS

PJ, HJ, JK, IYW and JKR designed the research. HJ, JK, and AHV performed cell experiments; YH and JKR designed the CMOS-MEA sensor device. PJ, YH, and JKR tested the CMOS-MEA sensor and wrote software. HJ, PJ, YH, JK, IYW, and JKR analyzed data. IYW and JKR acquired funding and supervised the research. HJ, PJ, YH, IYW and JKR wrote the manuscript with feedback from all authors.

