## Supporting Materials for "Label-Free All-Electrical Tracking of Individual and Collective Cell Migration on a Megapixel CMOS Capacitance Sensor"

### SUPPLEMENTARY MATERIAL

#### Note S1. CMOS capacitance sensor chip design

The capacitance sensor is a custom silicon chip fabricated in a commercial 180 nm CMOS process. Over the course of several years, we have designed several generations of related capacitance imaging chips [32–35, 51–54], and this new design has significantly more pixels, a larger active area, improved uniformity, and several new features.

The sensor has an array of 1024 × 1024 pixels, which are individually addressable through row/column control signals. Spatially-resolved capacitance measurements are made by rapidly scanning the address across the array to sequentially measure each pixel. To acquire a full frame 1024 × 1024 capacitance image takes less than 1 minute.

Figure S1a and b show an image of the sensor chip wirebonded to a small printed circuit board module and connected to a custom data acquisition board. The chip is controlled through an FPGA, which connects via USB to a Python environment.

Supplementary Table S1. Summary and performance comparison with prior literature

| Parameter | This Work | [32] | [36] | [9, 16] | [15? ] |
|---|---|---|---|---|---|
| **FOV** | 10.2 x 10.2 mm ($104.04\ mm^2$) | 2.56 x 5.12 mm ($13.1\ mm^2$) | 3.6 x 1.9 mm ($6.84\ mm^2$) | 1.26 x 1.26 mm ($1.59\ mm^2$) | 0.232 x 0.232 mm ($0.054\ mm^2$) |
| **Pixel Size** | 10 x 10 $\mu m^2$ | 10 x 10 $\mu m^2$ | 2.25 x 2.25 $\mu m^2$ | 20 x 20 $\mu m^2$ | 0.6 x 0.89 $\mu m^2$ |
| **# of pixels** | 1,048,576 | 131,072 | 368,000 | 4,096 | 65,536 |
| **Estimated # cells tracked*** | ∼12,600 | ∼1,575 | ∼822 | ∼191 | ∼7 |
| **Power** | 7.2 mW | 58.8 mW | - | 1.25 W | 15 mW |
| **Experimental Demo** | Epithelial/Mesenchymal cells, tissues, spheroids | Biofilm | Microparticles | Epithelial cells, Cancer cells | Microparticles, Cancer cells |

***** Calculated based on a 24-hour tracking period where the effective territory of a single cell is modeled as $A = 4\pi Dt$. Using a median diffusion coefficient ($D$) of 25 $\mu m^2/h$ over $t = 24$ hours, the explored area per cell is ∼7,540 $\mu m^2$. The total number of trackable cells ($N$) is derived by dividing the total FOV area by this territory, adjusted by a hexagonal packing density factor of 90.6% to account for non-overlapping circular boundaries.

*Capacitance sensing modes*

Several different capacitance sensing modes are supported by the custom CMOS sensor:

- **Self-capacitance.** In the simplest sensing mode, one pixel is rapidly charged and discharged by non-overlapping clocks $\varphi 1$ and $\varphi 2$, generating a sensing current. The current is routed to one of three integrator output channels, and is converted into an output voltage (Fig S2a), which is digitized by an external ADC. The self-capacitance is the sum of fringe field coupling to other nearby pixels, plus vertical coupling to the bulk electrolyte.

- **Self-capacitance with neighbor shielding.** Optionally, the sensor can also be configured to switch neighboring pixels in phase with the sensing pixel (Fig S2b), shielding the lateral fringe fields so that only the vertical fields remain [34]. The shielding mode tends to increase the capacitive contrast between materials above the sensor and moderately enhance the SNR, illustrated in Fig S5 for measurements of deionized water.

- **Mutual capacitance.** The chip can also be configured to measure the pairwise mutual-capacitance ($C_m$) between any two pixels in the array (Fig. S2c). The signal diversity available from overlapping sets of $C_m$ measurements can enable a variety of computational imaging techniques, including super-resolution image reconstruction [33] and 3-D capacitance tomography [35].

- **Mutual capacitance with virtual pixel grouping.** Additionally, contiguous groups of pixels in the array can operated in parallel as a larger 'virtual pixel.' The mutual capacitance ($C_m$) can then be measured between 'virtual pixels'. Increasing the size of a virtual pixel enhances the signal strength and extends the depth of detection, at the cost of spatial resolution. A virtual pixel of size 2×2 is illustrated in Fig. S2d. Virtual pixel groups of arbitrary rectangular sizes are supported.

The capacitance measurements can be performed with switching frequencies up to 50MHz. The electrochemical interfaces exhibit some frequency dependence, and at higher frequencies, there tends to be a reduction in absolute capacitance; however, there are advantages to working at MHz frequencies due to reduced electrochemical Debye screening and improved detection depth [15].

Each pixel in the array is addressable and configurable, as shown in Fig S1c, d, and Fig S6. Non-overlapping clocks are provided row-wise, and signals are measured column-wise. For example, in a self-capacitance measurement, one pixel at a time is selected by R1, C1, and OUT_EN, and the integrated switched-capacitor current is measured at the bottom of the column. For a virtual pixel mutual capacitance measurement, consider the extreme case in which both virtual pixels have a size of 1. Two pixels are selected with (R1,C1,STDBY) and (R2,C2,OUT_EN); they are provided with opposite clock phases, and only the second pixel's current is integrated and measured. For larger virtual-pixel sizes, all pixels belonging to the same virtual pixel are simply configured with the same control logic.

Column and row logic is included to shift the control signals by one row (or column) at a time, rastering the entire control pattern through the full array to capture a whole image. Each pixel also contains 1 local memory bit to hold the electrode at a constant potential, which can be used for electrode characterization, electroplating, or stimulation

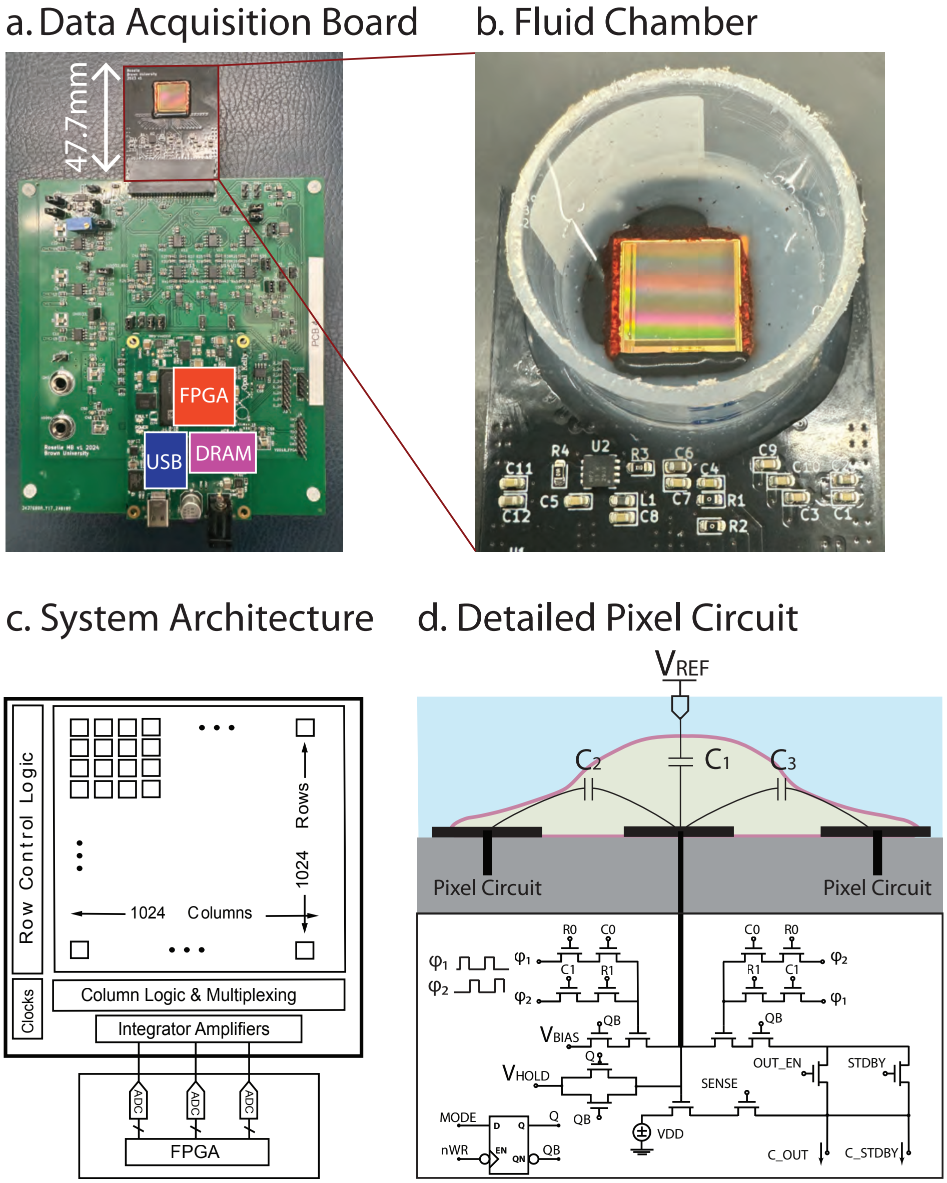


Supplementary Figure S1. (a) The chip is wirebonded to a small sensor module, which is connected to the data acquisition board with an FPGA on it. (b) Often, a simple open fluid chamber is mounted around the chip with silicone adhesive. (c) System architecture of the 1024 × 1024 sensor array. (d) Circuit schematic in each pixel. This is a switched capacitor circuit which is row-column addressable with non-overlapping clocks.

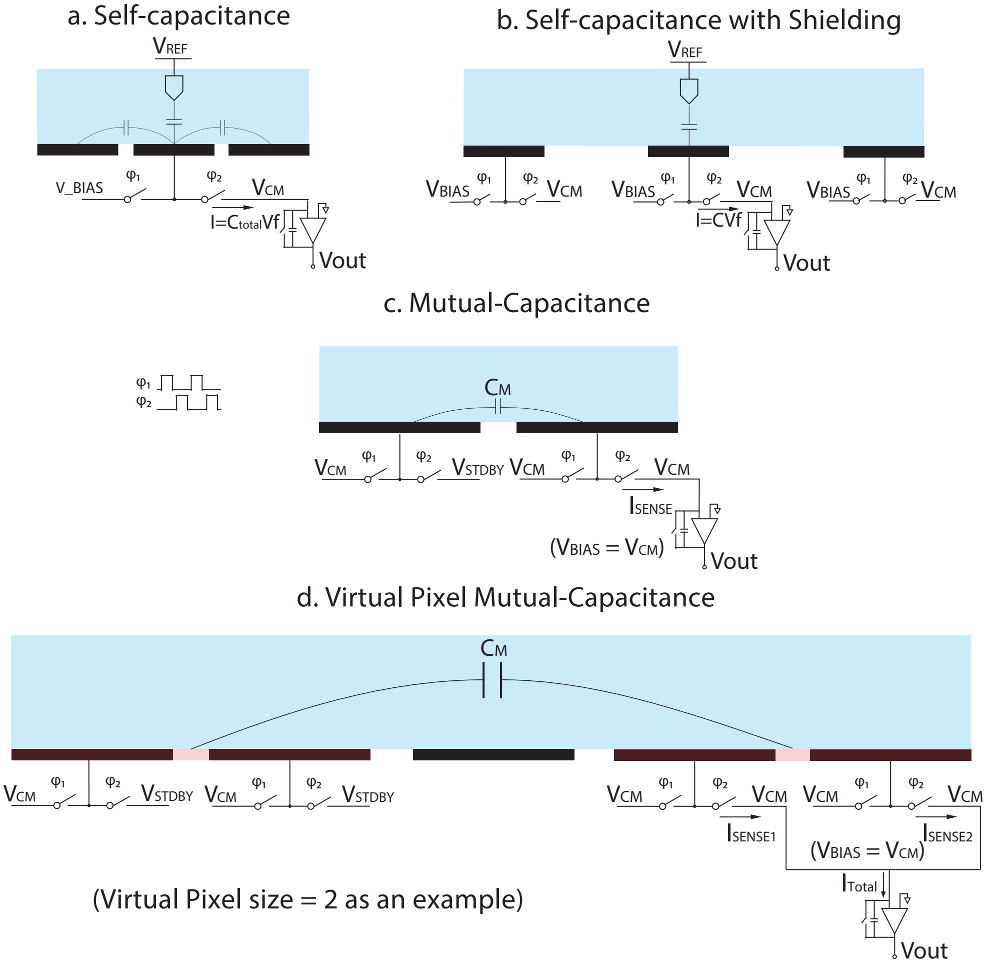


Supplementary Figure S2. (a) Circuit configuration in self-capacitance and (b) self-capacitance with shielding. The measured pixel with parasitic capacitance from surrounding pixels is shielded by aligning the switching clocks for all neighboring pixels. (c) Mutual-capacitance and (d) virtual pixel mutual-capacitance measurement (V = 2 as an example).

(Fig S1d). The chip provides analog outputs which are digitized by three external 500 kS/s 18-bit ADCs.

To help with calibration and testing, the pixels on the outer perimeter are dedicated for capacitance calibration, and contain four on-chip reference values. Fig S3 presents a histogram of these 4,092 calibration pixels, measured at 12.5 MHz. The observed standard deviation includes device variation and is generally less than 1 fF, but the true measurement noise floor can be as low as 0.1 attofards (rms), depending on the integration time. Measurements are performed serially through the array, and we usually operate at 24,000 pixels/sec, which translates to 5 $aF_{rms}$ electrical noise floor and less than 1 minute per frame, which is appropriate for time-lapse imaging in cell culture applications (Fig S4).

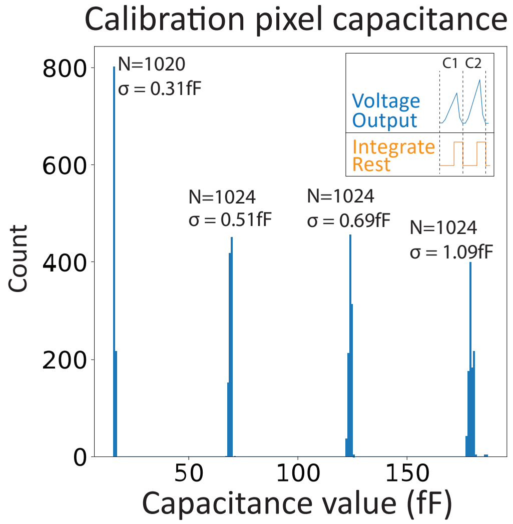


Supplementary Figure S3. Capacitance calibration. The on-chip calibration capacitors are used to calibrate the measurements, and are included in every measurement. Four test capacitor values are included around the outer perimeter of the chip. Shown here is a histogram of the calibration pixels from one image.

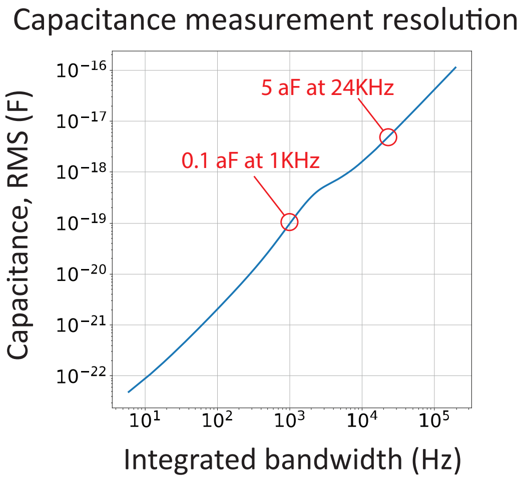


Supplementary Figure S4. Integrated measurement noise, expressed as RMS capacitance, as a function of the measurement bandwidth. The bandwidth determines the number of pixels acquired per second, and thus the frame rate.

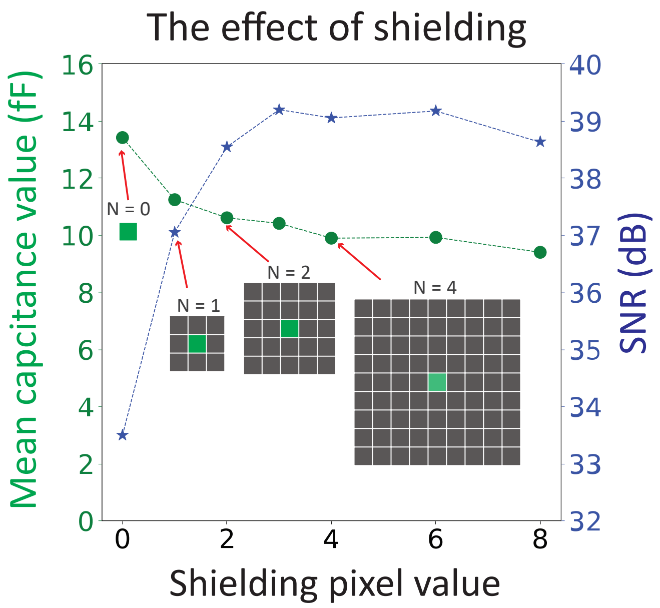


Supplementary Figure S5. Self-capacitance with shielding. Neighboring pixels can be switched in phase with the sensing pixel, to eliminate the lateral coupling in favor of the more vertical electric fields. As more of the neighboring pixels are included, the absolute capacitance is reduced but the signal contrast typically improves. Here capacitance was measured in uniform DI water. The SNR is defined as the ratio of the mean measured capacitance to its standard deviation.

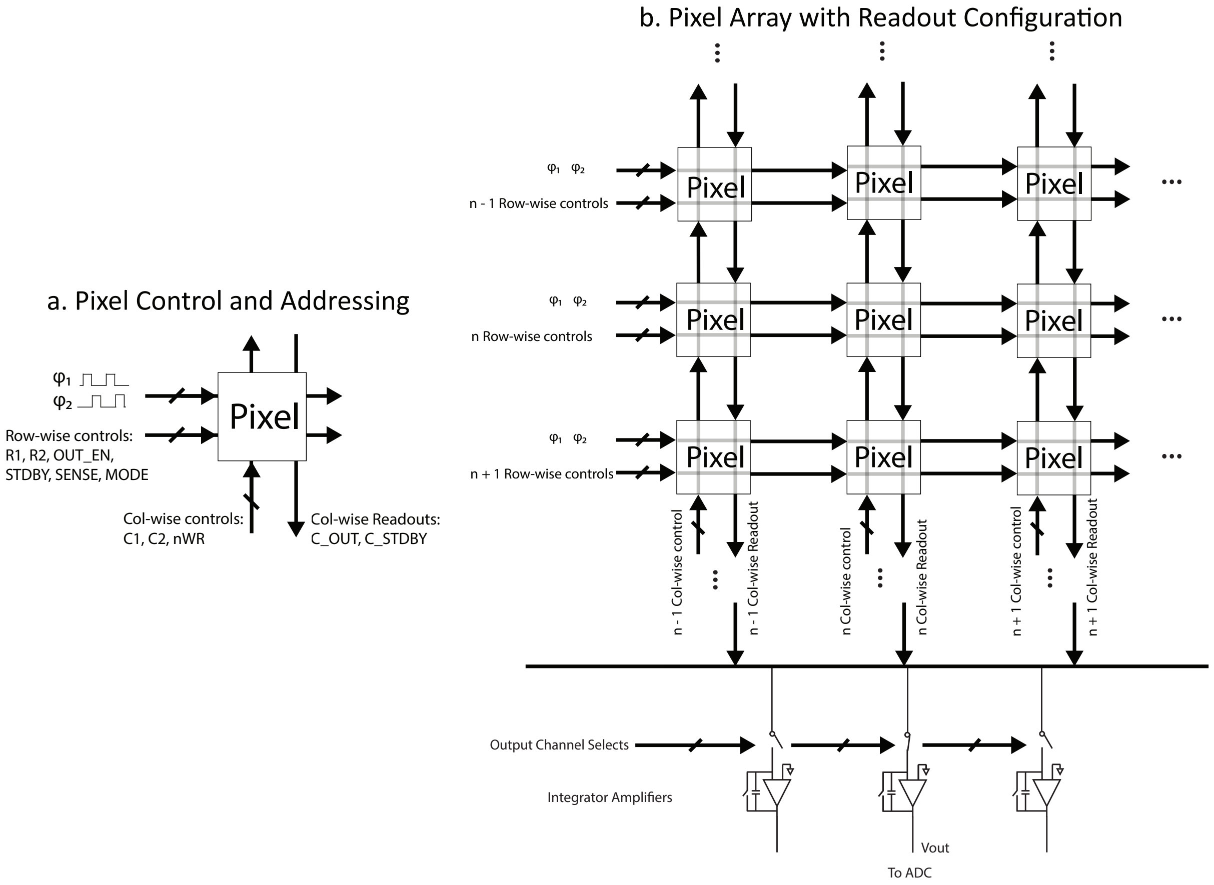


Supplementary Figure S6. (a) The pixels are addressable switched-capacitor circuits using row-column addressing and non-overlapping row-wise clocks. (b) An array of pixels with the readout configuration (CH2 is selected as the output channel, as an example).

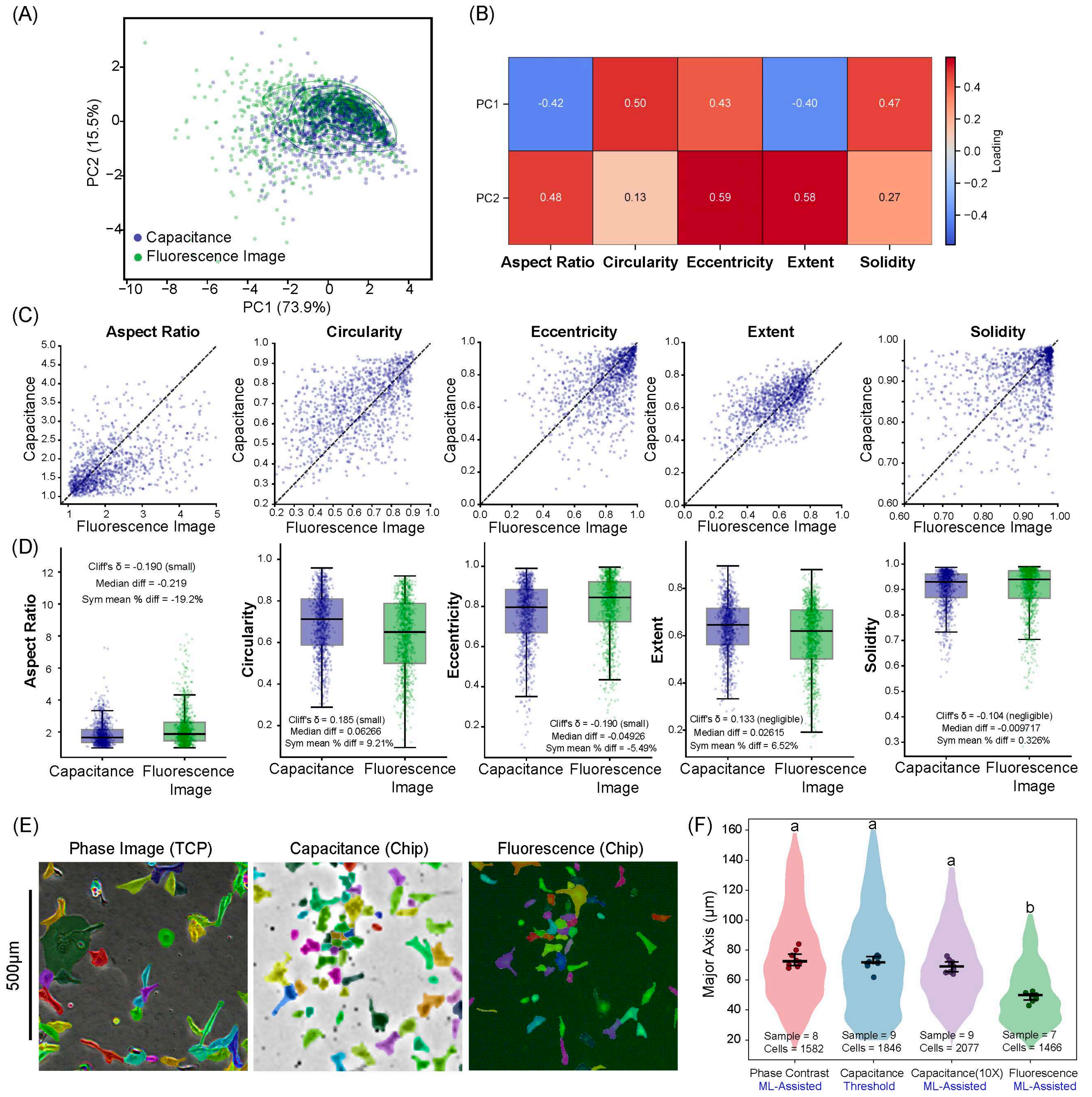


Supplementary Figure S7. **Comparison of cell morphology obtained from different signal acquisition methods.** (A) Distribution of cells in principal component (PC) space based on capacitance and fluorescence measurements on the CMOS MEA. Contour lines represent five KDE-based density levels. (B) Principal component loadings for each morphological feature. (C, D) Comparison of morphological features derived from capacitance and fluorescence intensity measurements. Each dot in (C) represents a matched pair obtained from the two methods, and the dotted line indicates the y = x reference line. (E) Representative snapshots of cell masks obtained from different imaging modalities: phase-contrast imaging on a tissue culture plate (TCP), capacitance imaging on a CMOS MEA, and fluorescence imaging on a CMOS MEA. Scale bar = 500 $\mu$m. (F) Comparison of cell major-axis length measured using different imaging methods and algorithm, CMOS images without 10× upscaling were segmented using threshold-based methods, whereas CMOS images with 10× upscaling were segmented using ML-assisted segmentation. Violin plots show cell-level distributions, while statistical comparisons were conducted using sample medians, using Welch's ANOVA followed by pairwise Welch's t-tests with Holm correction for multiple comparisons. Different letters indicate statistically significant differences(adjusted p ¡ 0.001) between groups; groups sharing the same letter are not significantly different.

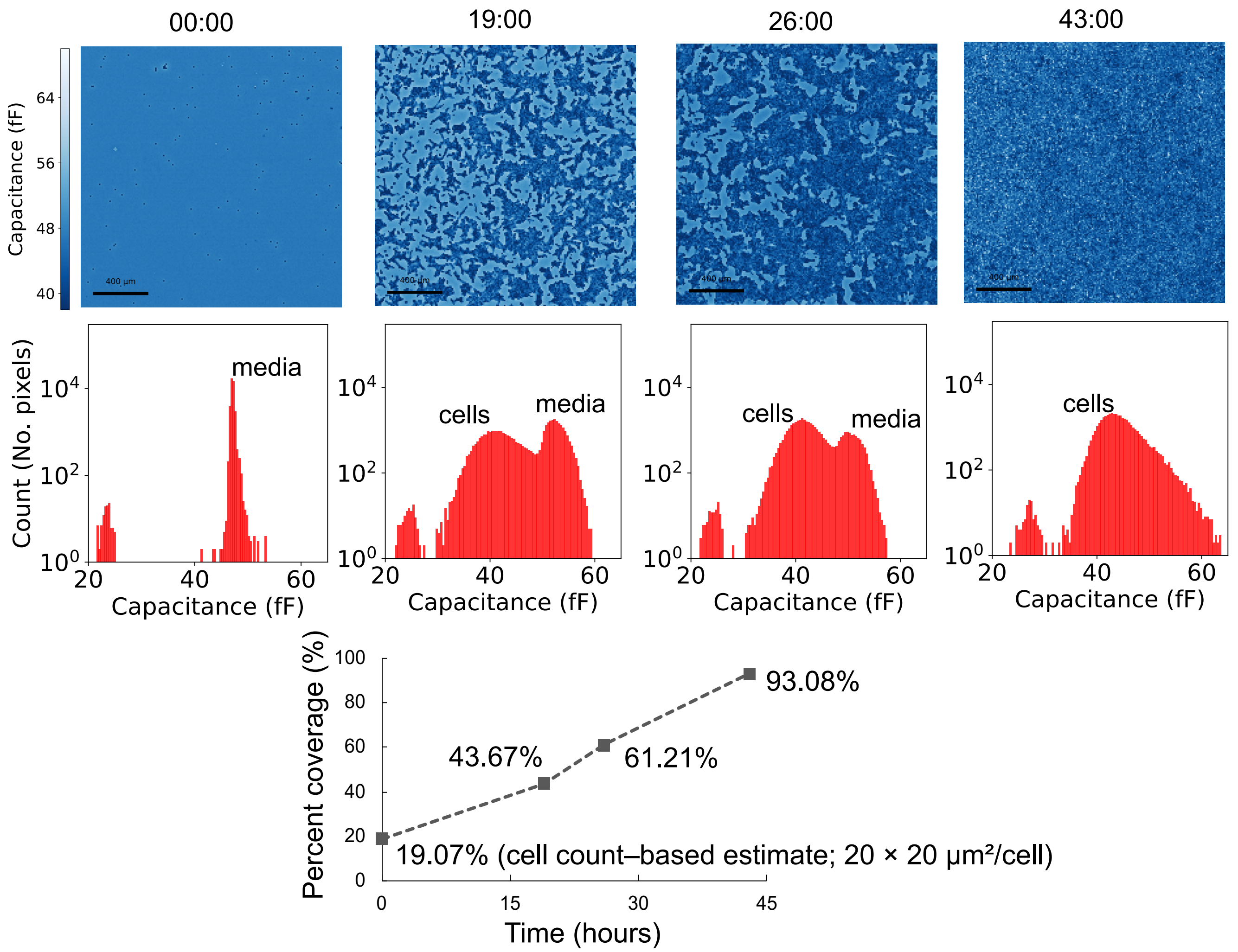


Supplementary Figure S8. Capacitance measurements were performed on non-transformed breast epithelial cells (MCF-10A) cultured on the CMOS chip. The cells gradually increased in density and reached confluence within approximately two days. By comparing cell images acquired at 19 hours(43.67% of coverage) and 43 hours after seeding(93.08% of coverage), we observed a substantial increase in cell coverage over this period, suggesting a doubling time of less than 24 hours. The initial coverage value was estimated from the cell number($5 \times 10^4$ cells) by assuming an average projected cell area of $20 \times 20 \mu m^2$ per cell, because the cell covered area at the initial time point could not be directly segmented.

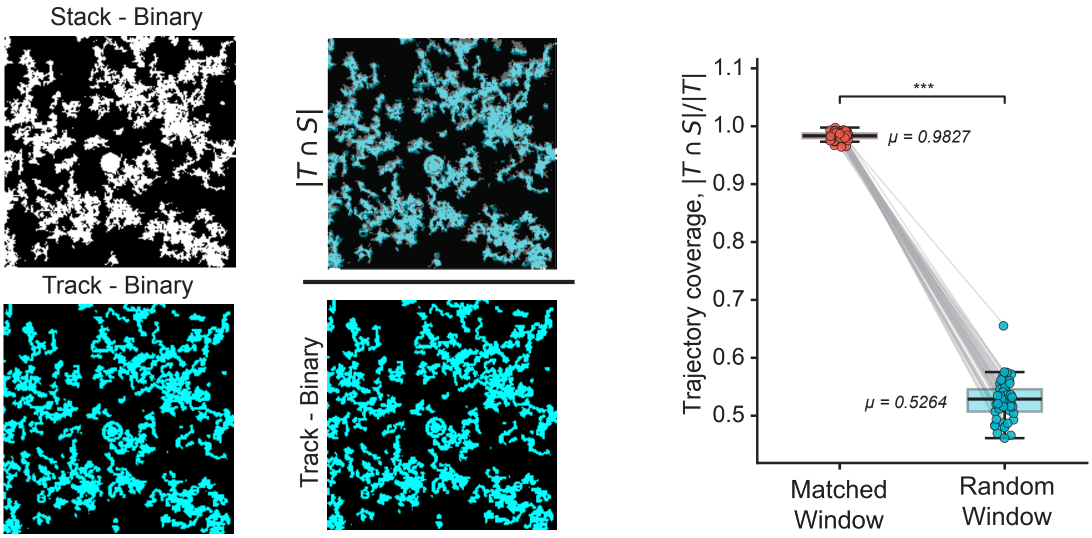


Supplementary Figure S9. **Verification of the single cell tracking method by comparing the overlap between cell image projection and trajectories.**

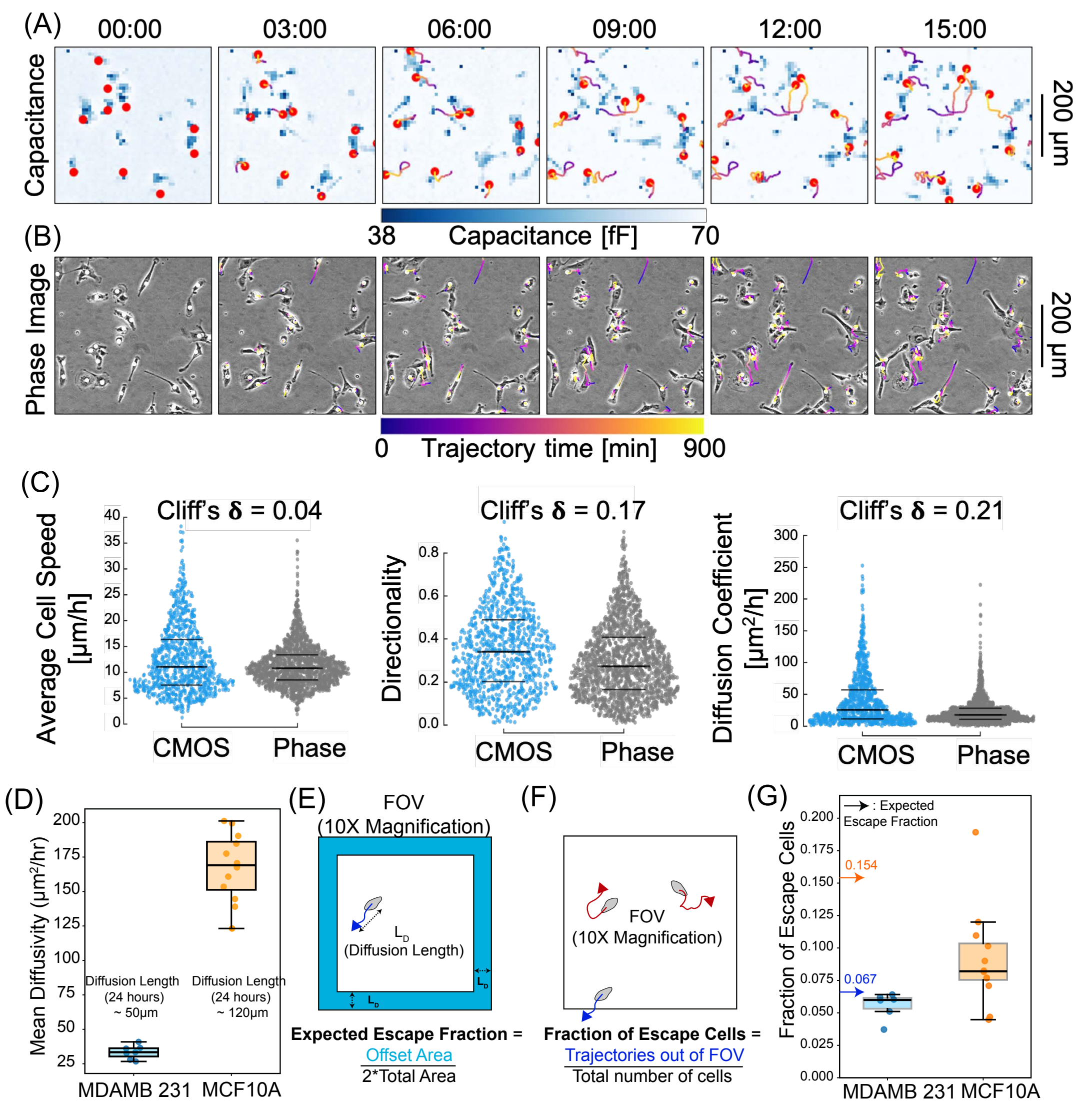


Supplementary Figure S10. **Effect of limited field of view on cell migration tracking due to cell escape.** (A) Cell migration on CMOS chip visualized with capacitance measurement. (B) Cell migration on a standardized TCP captured with phase contrast imaging. (C) Quantitative comparison of migratory phenotypes between cells on the CMOS and on TCP substrate. Average cell speed $= \frac{\text{Distance}_{\text{Total}}}{\text{Time}_{\text{Total}}}$, and, Directionality $= \frac{\text{Displacement}_{\text{Total}}}{\text{Distance}_{\text{Total}}}$. Each dot on the scatter plots represents data from a single cell. The Cliff's $\delta$ values suggest negligible or small deviations in these parameters. (D) Estimated cell migration diffusivity for MDAMB-231 and MCF10A cells. (E) Schematic of the expected escape fraction calculation. The boundary-associated escape region was defined using half of the diffusion-length offset from the FOV edge. (F) Schematic of escape-cell identification from trajectory data. Cells were classified as escaping when their trajectories terminated or became arrested within <5 pixels of the image boundary. (G) Comparison of the expected escape fraction and the experimentally measured escape fraction.

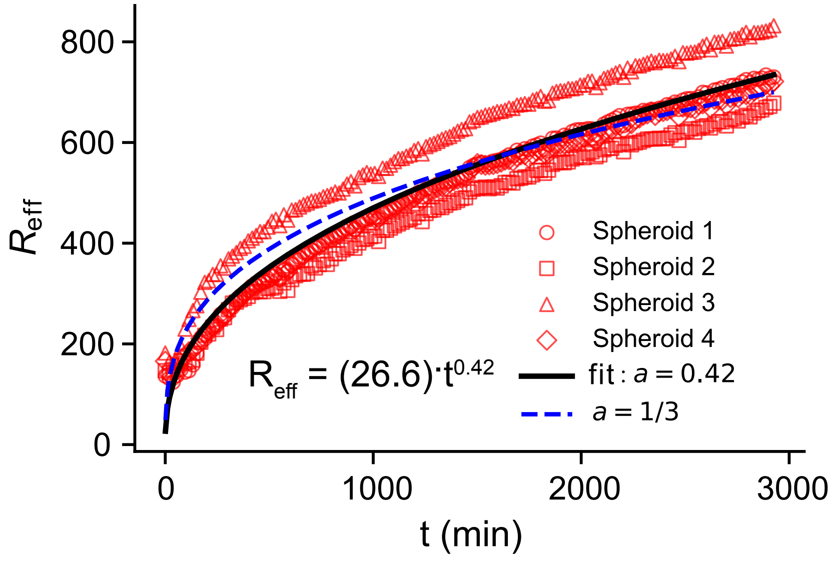


Supplementary Figure S11. Spheroid spreading tracked over time on a CMOS chip. The two-dimensional spreading area was quantified over time from (N = 4) spheroids. The blue dotted line indicates a earlier active-wetting fit with a power-law exponent of 0.3, whereas the black solid line shows the fit to the experimental data.

Figure S12: Virtual Pixel Detection Depth COMSOL Simulation

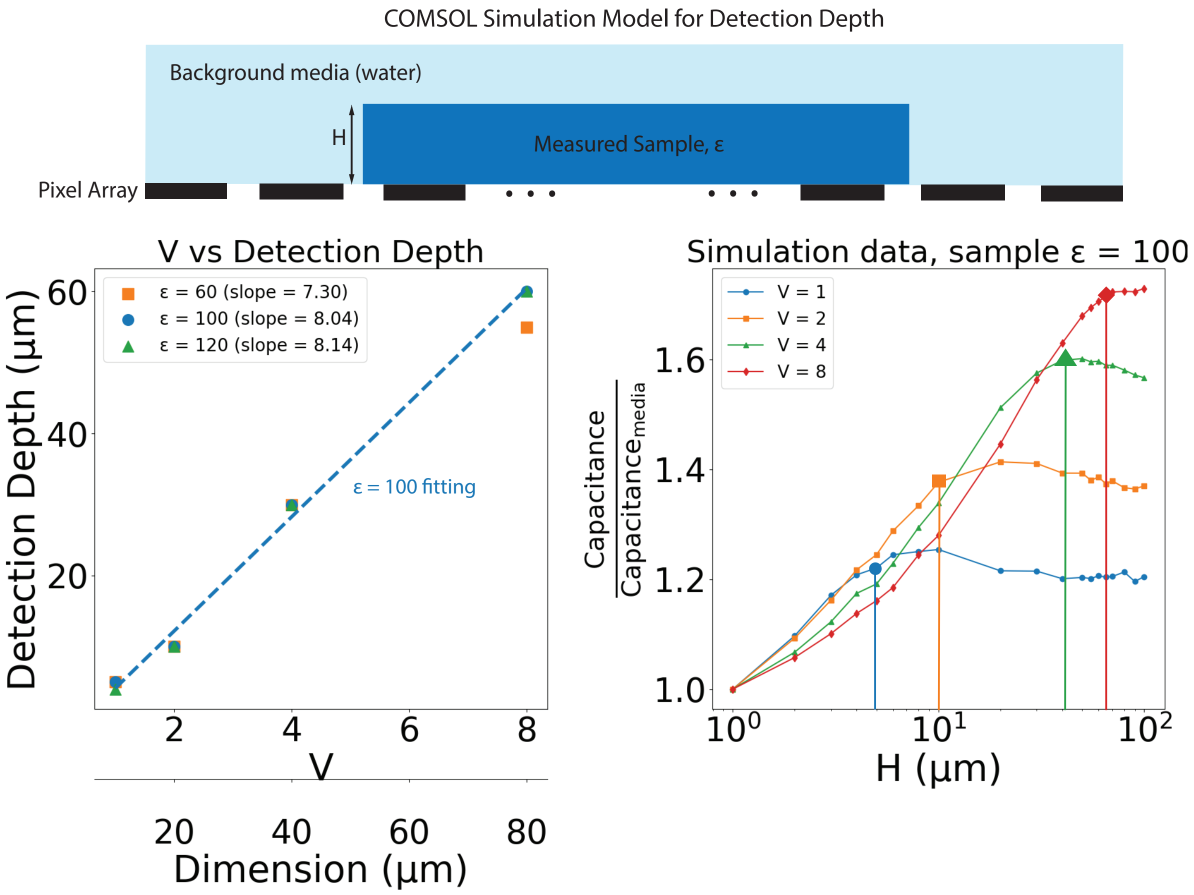


Supplementary Figure S12. Detection depth obtained for varying sizes of virtual pixel using CMOSOL simulation by updating the height (H) of the measured sample, until the capacitance value begins to saturate. The detection depth remains almost constant even when the dielectric constant $\epsilon$ changes.

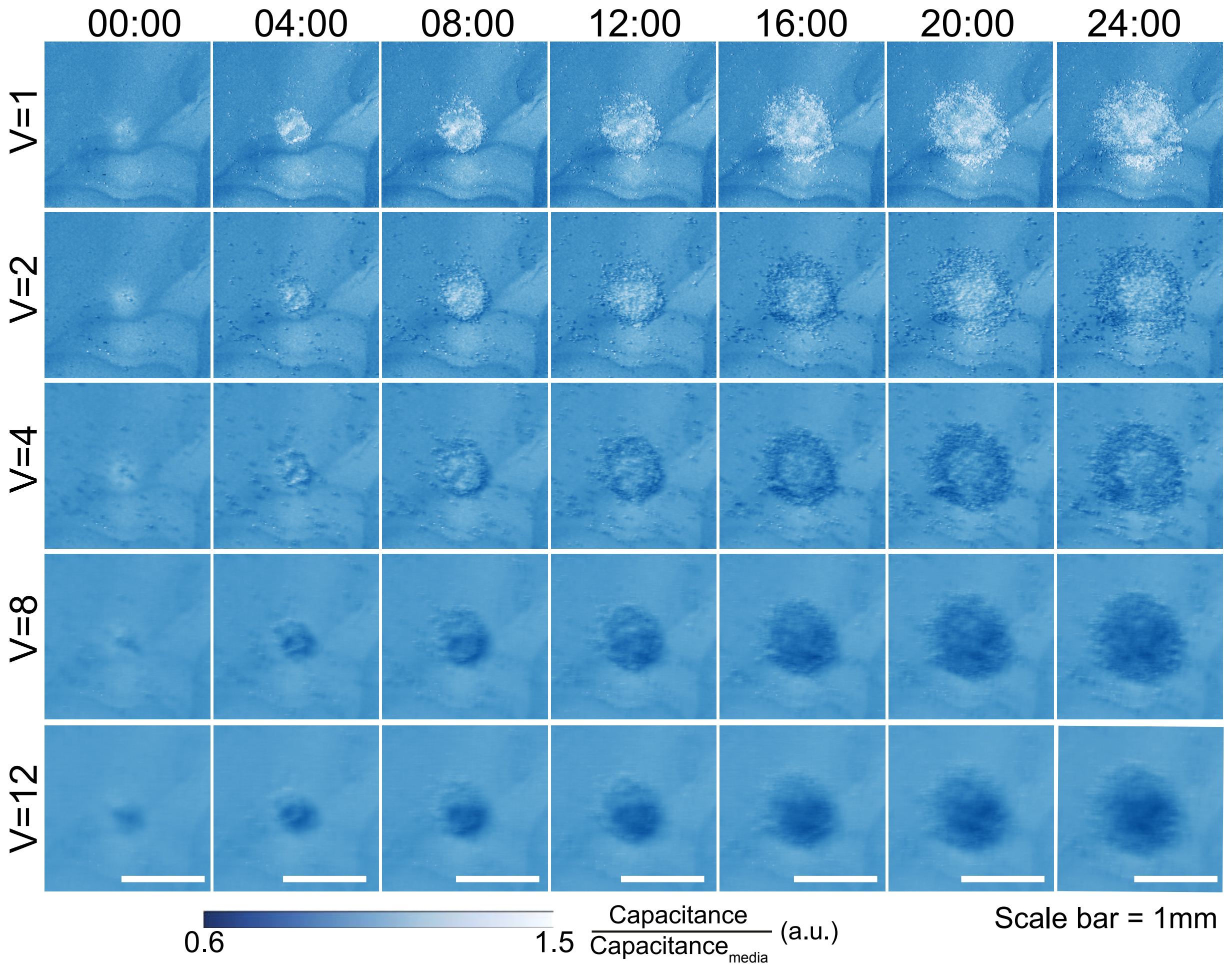


Supplementary Figure S13. Normalized capacitance measurements obtained using different virtual-pixel sizes(V=1,2,4,and 8) in mutual capacitance imaging of representative spreading spheroid.

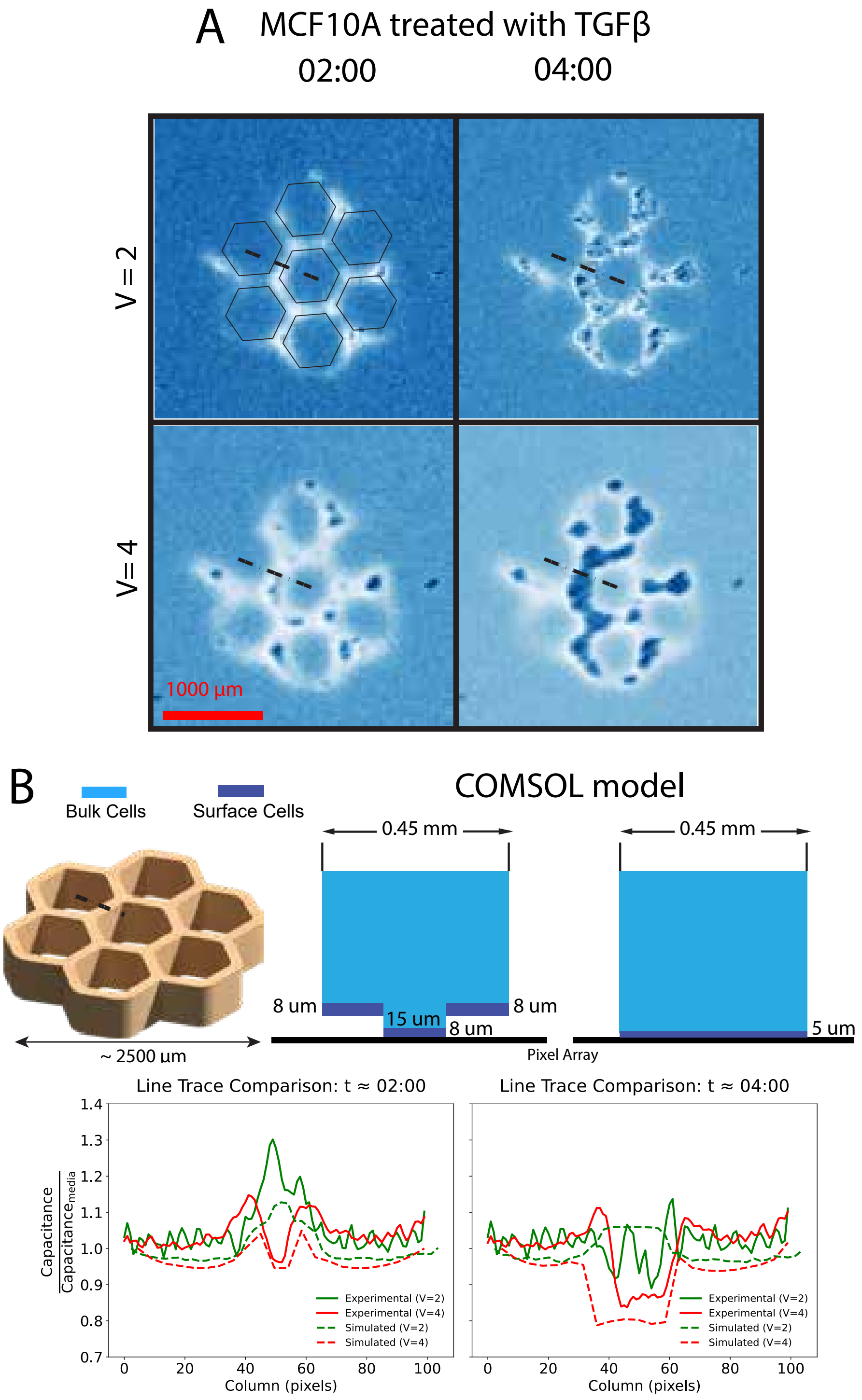


Supplementary Figure S14. Monitoring MCF10A treated with TGF$\beta$ honeycomb tissue after depositing on the CMOS sensor. (A) Virtual pixel images with V = 2, 4 at early time points. (B) The simplified COMSOL model, along with a comparison of simulation and experimental results for virtual pixel sizes of 2 and 4 at 2 h and 4 h, in the cross-sectional region indicating the dynamics of tissue contact on the sensor.
